# Constitutive Androstane Receptor contributes towards increased drug clearance in cholestasis

**DOI:** 10.1101/520692

**Authors:** Bhoomika Mathur, Waqar Arif, Megan Patton, Rahiman Faiyaz, Auinash Kalsotra, Antony M Wheatley, Sayeepriyadarshini Anakk

## Abstract

Understanding the alterations in drug metabolism in different liver diseases is crucial for appropriate therapeutic intervention. We performed high-throughput RNA sequencing on various liver injury models, including cholestasis, diet-induced steatosis, and regeneration. Comparative liver transcriptome analysis revealed overlapping and distinct gene profiles among different liver diseases. Particularly, cholestatic livers displayed robust induction of drug metabolizing genes. This upregulation is not a generic hepatic stress response, as it was suppressed or unchanged in other models of liver diseases. Consistently, drug metabolic gene profiles were induced in a subset of biliary atresia patients, but not in individuals with hepatitis B or C viral infection, and alcoholic hepatitis. Further analysis revealed this induction was specific to genes regulated by nuclear receptor CAR (Constitutive Androstane Receptor). To test this, we challenged cholestatic mice with a paralytic agent, zoxazolamine. Compared to controls, these mice displayed significantly reduced paralysis time, reflecting increased drug metabolism, and this effect was lost upon inhibition of CAR. Thus, CAR activation can alter therapeutic efficacy of certain drugs in a subset of cholestatic individuals.

## Introduction

The liver plays a fundamental role in drug metabolism and detoxification. It is known that multiple factors, including genetics, diet, alcohol and viral infections can compromise its function. For instance, cholestatic liver disease can be drug-induced or secondary to mutations or due to immune modulation (1, 2) (3). On the other hand, high-fat and western-diet primarily drive the development of fatty liver disease, while excess alcohol consumption, hepatitis B and C infections can lead to cirrhosis and eventually cancer. As different therapeutic interventions are employed to treat these conditions, it is important to understand the molecular networks that are perturbed during these disease states.

Here, we used unbiased high throughput approach and performed comparative analysis of transcriptome of different liver diseases to identify the common and distinct gene signatures between them. RNA-sequencing data from mouse models of cholestasis, diet-induced fatty liver disease and liver regeneration revealed alterations in genes involved in inflammation, proliferation, and lipid metabolism. Intriguingly, CYP450 pathway was specifically upregulated in cholestatic livers, whereas it was downregulated in steatosis and regeneration. To examine if this expression pattern is maintained in humans, we analyzed publicly available liver transcriptome datasets of Hepatitis B and C viral infections (4), alcoholic hepatitis (5), and biliary atresia (6). We did not find upregulation of P450s in any of the conditions except for a subset of biliary atresia patients with extremely high GGT levels.

This is surprising since liver function is compromised during cholestasis, one would anticipate reduced metabolic capacity. Such confounding data have been reported, in the context of Primary Biliary Cirrhosis (PBC), liver cirrhosis, and biliary obstruction (7–9). It is important to note that some of the phase I metabolic genes such as *Cyp2bs* and *Cyp3as* can hydroxylate and facilitate bile acid (BA) excretion (10) and increased drug clearance during cholestatic liver diseases have been previously noted (11, 12).

The major regulators for phase I xenobiotic metabolism are nuclear receptors Constitutive Androstane Receptor (CAR) and Pregnane X Receptor (PXR) (13). Therefore, we examined the Phase I genes in cholestasis mouse model and found a significant correlation with CAR activation, while Phase II genes correlate with activation of both CAR and PXR. We also determined that the observed increase in detoxification machinery is associated with the BA burden. Taken together, our findings reveal that CAR activation in cholestasis may occur to promote BA excretion but leads to unintended consequence of increased drug clearance.

## Results

### Transcriptomic profiling of diseased murine livers revealed upregulation of genes involved in metabolism during cholestasis

Liver is a central hub responsible for the breakdown of carbohydrates, fat, protein and foreign compounds including prescription drugs. To examine how transcriptional liver networks are altered during different liver injuries, we performed and analyzed RNA-seq on cholestatic, fatty and regenerating mouse livers. Global deletion of both Farnesoid X Receptor (*Fxr*) and Small Heterodimer Partner (*Shp*) (DKO) resulted in cholestasis, which is excessive accumulation of bile acids (BA) (14) (15). Histological analysis of DKO livers showed liver injury such as hepatocyte ballooning, ductular reactions, and focal inflammation, consistent with severe cholestasis. Further, high fat diet fed mice developed macro-steatosis and accumulated large lipid droplets in the liver and partial hepatectomy resulted in a robust regenerative response, which is indicated by the presence of mitotic bodies (Supplementary Fig. 1). We found overlapping and distinct gene programs switched on in the livers after these different conditions (Fig. 1A). Gene ontology analysis of overlapping pathways specifically enriched the cellular proliferative response (Supplementary Table 1). Interestingly, many metabolic pathways were up regulated, while DNA damage, hypoxic and insulin response were down regulated during cholestasis (Fig. 1B). On the other hand, high fat diet-fed mice as expected showed upregulation of genes related to lipid transport, immune response activation, and acyl coA metabolic pathways. Downregulated genes were related to glucose homeostasis, oxidation-reduction, steroid, and P450-dependent metabolic pathways (Fig. 1C). Regenerating livers showed upregulation of genes related to cell cycle, mitosis, and acute phase response, and downregulation of oxidation-reduction, steroid, glucose, and P450-dependent metabolic pathways (Fig. 1D).

**Fig. 1.**
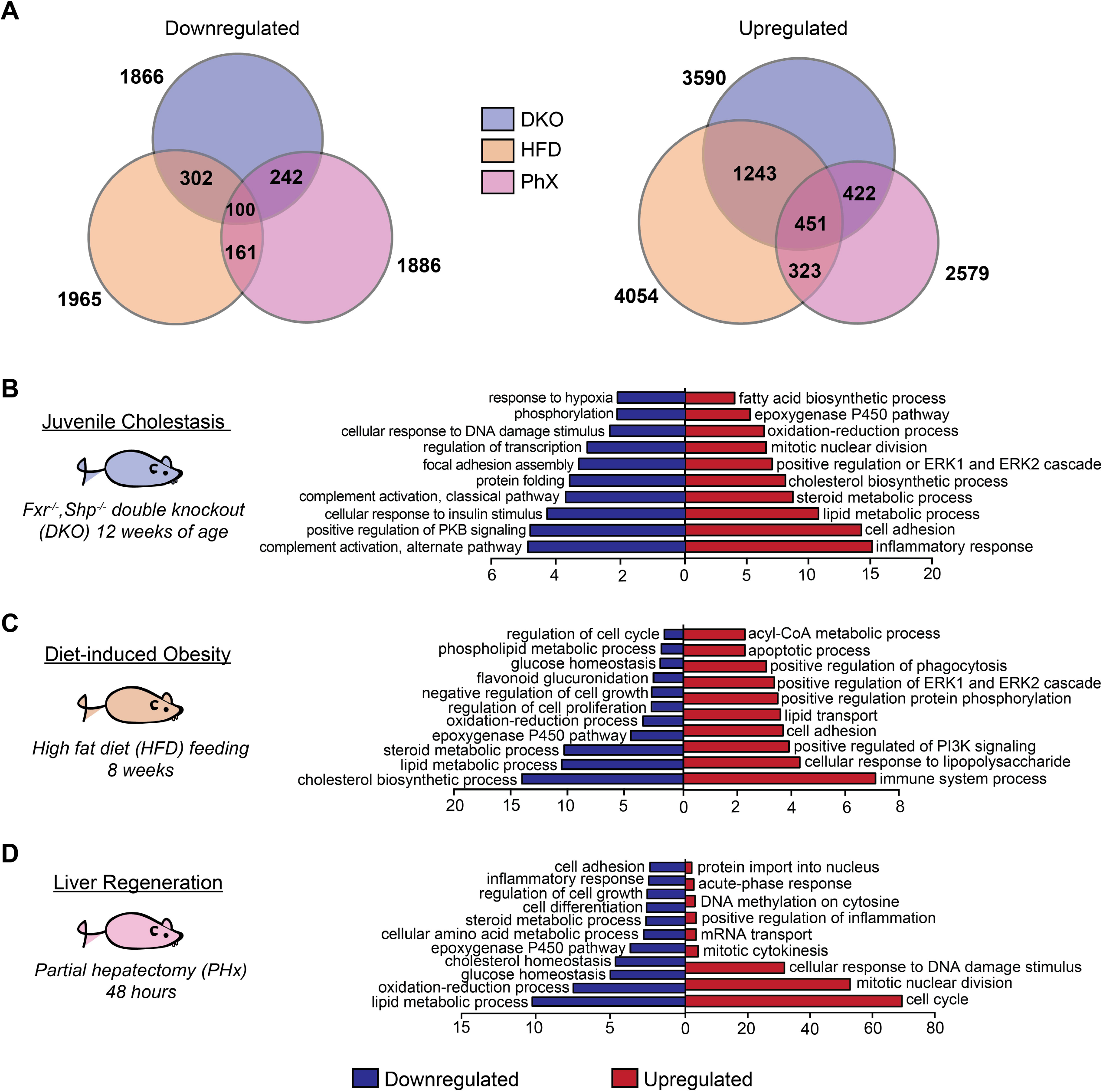
Genome wide profiling reveals common and unique gene networks in cholestasis, steatosis and regeneration. (**A**) Venn Diagrams represent the overlap of downregulated and upregulated gene expression among cholestasis (DKO), diet-induced steatosis (HFD) and regeneration (PHx). DAVID Gene ontology pathway analysis was performed on upregulated and downregulated genes in (**B**) cholestatic, (**C**) fatty, and (**D**) regenerating livers (n=2-3 mice per group, Fold change >1.5, FDR <20).

Thus, an increase in metabolic gene profile of the liver is not a general liver injury response but is rather specific to cholestasis. We focused our interest on drug metabolism and examined the transcriptome of CA-fed WT mice livers. In addition to induction of cell cycle and mitosis related genes, CA diet caused pronounced upregulation of oxidation-reduction, glutathione metabolism, and cellular drug response pathways (Supplementary Fig. 2A). These results demonstrate that BA overload can activate cytochrome P450s, and subsequently, drug metabolizing phase I and phase II genes as observed in DKO mice (Supplementary Fig. 2B-C). We examined the circulating BA composition between CA-fed control and DKO mice (Supplementary Fig. 2D) and found increases in both 6α (all muricholates) and 12α hydroxy BAs (both DCA and CA) as previously reported (14) (16).

To ascertain if phase I metabolism is induced in human liver diseases, we analyzed the liver transcriptome post-HBV or -HCV infection (4) to determine if the drug metabolic changes are secondary to viral infection and/or inflammation. In this group, we separated the samples into two cohorts based on whether the disease progressed to hepatocellular carcinoma (cHBV, cHCV) or not (HBV, HCV). Additionally, we mined the transcriptomic data from patients with severe alcoholic hepatitis (5). The transcript changes displayed overlap between HBV and HCV cohorts but were distinct from alcoholic hepatitis group (Fig. 2A). We did not find any alteration in the detoxification machinery (Fig. 2B, C) post HBV or HCV infection, whereas in alcoholic hepatitis, down regulation of phase I and II genes, except *Cyp1B1* and *Sult1C2* was observed (Fig. 2B, C). Also, many of the *Cyp* genes were significantly decreased in cHBV group (Fig. 2C), while specific downregulation of *Cyp2B6, 2C19* and *3A4* was observed in cHCV cohort (Fig. 2B). On the other hand, Phase II genes, *Sult1C2* and *Ugt1A6*, were upregulated in both cancer groups (Fig. 2C).

**Fig. 2.**
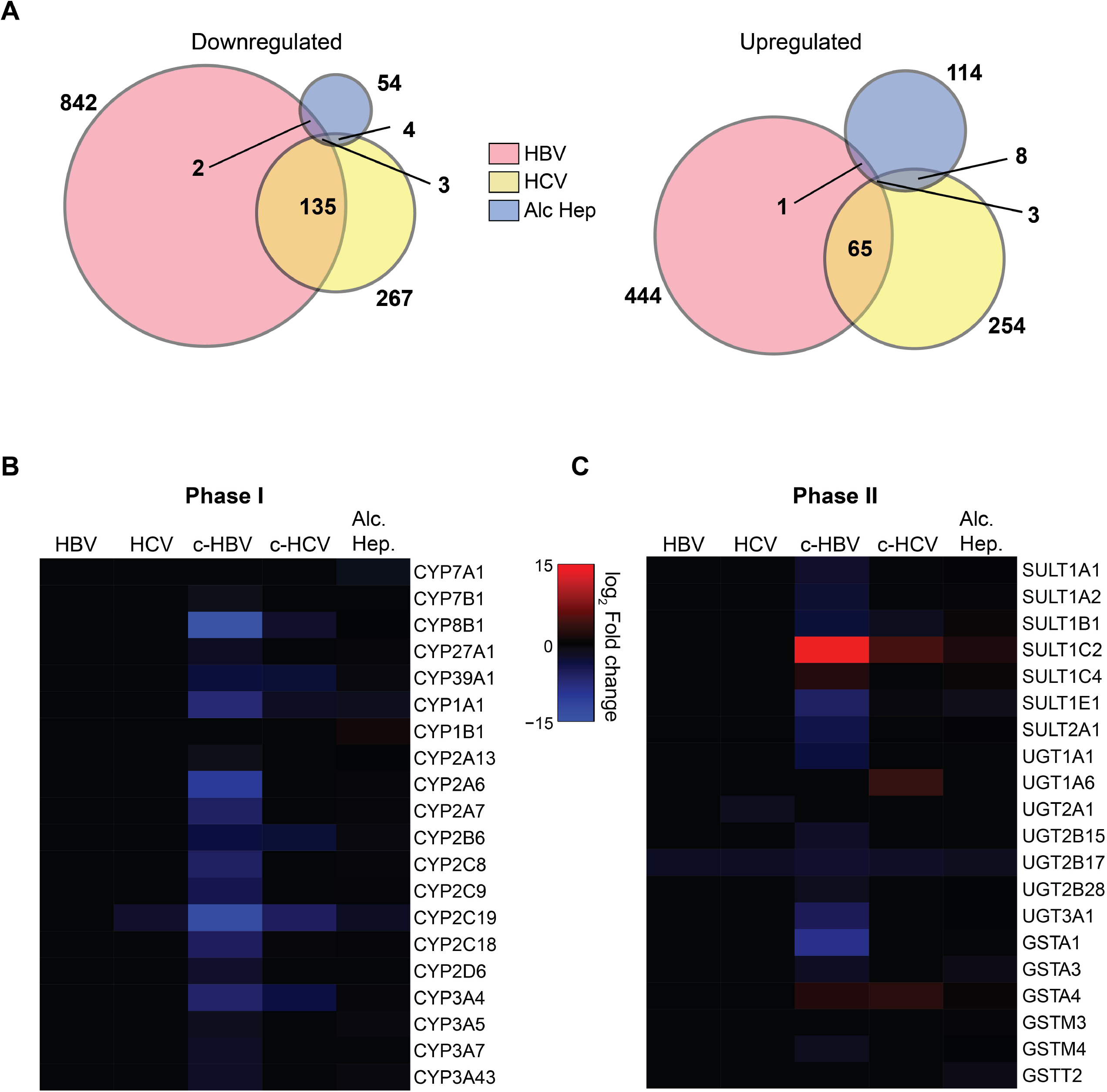
Comparative transcriptomic analyses indicate that drug metabolism is not a generalized response to liver injury. Previously conducted microarray data analyses for liver injury models in patients like Hepatitis B and C, cancer associated HepB (c-HBV) and HepC (c-HCV), Alcoholic Hepatitis (GEO dataset: GSE28619) were mined for alterations in gene expression. (**A**) Venn diagrams show negligible overlap in gene expression among these liver diseases. Heat maps display no induction, but rather downregulation of (**B**) Phase I and (**C**) II drug metabolic genes in these diseases (Fold Change > 1.5, p value <0.05).

Furthermore, cholestasis subsequent to deletion of *Bsep*, a major BA canalicular exporter, did not activate, but suppressed, phase I metabolism (17). Notably, their overall serum BA levels were dramatically lower compared to DKO mice (15), suggesting that a particularly high threshold of BA concentration in hepatocytes may promote detoxification pathways such as Phase I and II, as a fail-safe mechanism to hydroxylate and excrete BAs.

### A common overlap is observed between DKO livers and CAR/PXR activation gene signatures

It is well known that nuclear receptors Constitutive Androstane Receptor (CAR) and Pregnane X Receptor (PXR) play a crucial role in the regulation of detoxification machinery (18, 19). Therefore we performed transcriptome analysis of livers after activation of CAR (TC treated) and compared the transcriptome with the previously published PXR-activated (PCN treated) data set (20) (Fig. 3A, B) to identify which of these two programs were contributing towards the increased expression of drug metabolizing genes in the cholestatic DKO livers. As expected TC and PCN treatment induced pathways associated with oxidation-reduction, xenobiotic, and steroid metabolism (Fig. 3A and B). We also found a strong upregulation of cell cycle associated transcripts with TC, but not with PCN treatment suggesting that CAR may play an important role in controlling liver proliferation. 12-week-old DKO livers revealed an 85% (11/13 genes) overlap with the CAR activated, and 37.5% (3/8 genes) overlap with the PXR activated, phase I gene signatures (Fig. 3C). We found that the overall transcript changes in DKO livers displayed a 37% overlap with CAR activation (Supplementary Fig. 3A). Moreover, we examined and found 61% of Phase I and 28% of Phase II gene targets altered in the DKO mice overlapped with that of TC-treated WT mice (Supplementary Fig. 3B-C).

**Fig. 3.**
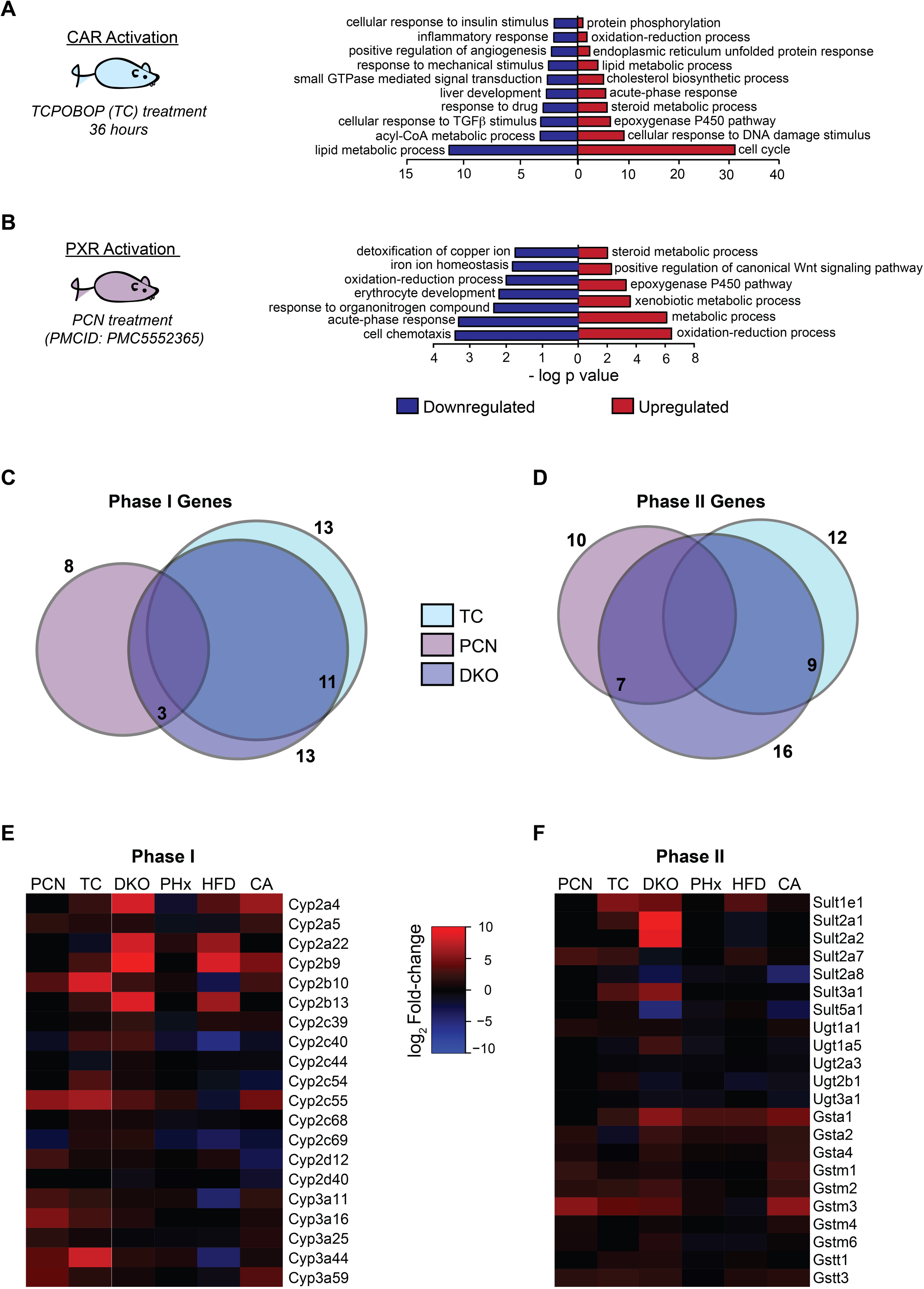
CAR target signatures are induced specifically in cholestatic conditions. Bar graphs show upregulated and downregulated biological pathways in (**A**) CAR-, and (**B**) PXR-activated livers (n=2-3 mice per group, Fold change >1.5, FDR <20). Venn diagrams show major overlap of (**C**) Phase I genes between TC-treated and DKO mice. (**D**) Phase II genes in DKO livers are equally spaced between TC- and PCN-treated mice. Numbers indicate total number of genes in individual conditions, and overlap of DKO with TC and PCN (n=3 per group, Fold change > 1.5, p value <0.05). Heat maps represent that the cholestatic DKO livers recapitulate profiles of (**E**) Phase I and (**F**) Phase II genes observed in TC treated (CAR activated) livers. Importantly, some of these gene changes were mimicked in bile acid (CA) diet, but not in PCN-treated (PXR activated), High Fat diet (HFD), and partial hepatectomized (PHx) livers.

We then examined individual gene expression of Phase I genes and compared their expression pattern across the different cohorts (Fig. 3E). As expected, *Cyp2bs*, and *2cs* were more responsive to CAR, while *Cyp3as* were more responsive to PXR (Fig. 3E). CA-fed livers displayed increases in Phase I genes, mimicking the profile of cholestatic DKOs. Compared to our previous microarray-based analysis of 5-week old DKO livers (14), RNA-seq results from 12-week old DKO mice showed broader, and more robust differences in gene expression (Fig. 3E). This indicates that worsening of cholestatic injury with age triggers a stronger and more extensive compensatory response. Further, forty-eight hours post partial hepatectomy (PHx), all the *Cyp* genes remained suppressed, consistent with reduced metabolic function during regeneration (21). Many cytochrome P450 genes were downregulated in diet-induced steatosis, except for *Cyp2b9, Cyp2b13, Cyp2a4* and *Cyp2a22* (Fig. 3E) as previously described (22). These increases in *Cyps* could be attributed to glucocorticoid receptor signaling, which is enhanced in steatosis (23).

Phase II sulfotransferase (*Sult*) genes were robustly induced only in DKO and TC-treated WT mice. Whereas, increases in glutathione sulfotransferases (*Gsts*) were observed in DKO, CA-fed, PCN- and TC-treated mice. None of the *Gsts* were increased in the steatotic and regenerating livers, except *Gsta1* (Fig. 3F). The expression of phase II genes in DKOs was similar between activation of either PXR (7/16) or CAR (9/16) (Fig. 3D), indicating that both receptors contribute to this process in DKO livers.

### Fxr and Shp network can interact with the detoxification pathway

FXR can induce PXR (24), and alter *Cyp3A4* expression in humans (25), therefore we examined the consequences of *Fxr* or *Shp* deletion on detoxification machinery. We compared the CAR-PXR signatures between DKO, FXR knockout (FXRKO), and SHP knockout (SHPKO) livers. *Cyp2b* and *Cyp2c* gene expression profiles in FXRKO and SHPKO did not match that of DKO livers. However, *Cyp2a5* expression showed dependency on FXR (Supplementary Fig. 4A, B). Interestingly, 85% (6/7) of Phase II genes induced in DKO overlapped with FXRKO except for *Sult2a1* (Supplementary Fig. 4C, D). On the contrary, SHPKO livers had lower *Gsta1, 2* and *Gstm2, 3*. Next, we evaluated a role for human *Fxr* and *Shp* in xenobiotic metabolism, since we know that expression of both these nuclear receptors are reduced during biliary atresia (26). We used previously published data (6) and compared biliary atresia samples with each other (22-169 days old) rather than comparing with control deceased liver samples (1.8 - 3.5 years old), since *Cyp* expression changes markedly with age (27). We found that xeno-sensing gene expression pattern segregated into three categories based on gamma-glutamyl transferase (GGT) levels - low (<450 U/L; n=21), mid (450-850 U/L; n= 31), or high (>850 U/L; n=17) (Fig. 4A, B, Supplementary Fig. 5A, B). It is important to note that out of the 69 biliary atresia samples, about 25% fall into the high GGT category. Interestingly, the expression profiles of several phase I genes including *Cyp2C8* and *Cyp3A4* genes were induced in high GGT samples (Fig. 4A). However, the high coefficient of variation (97.40%) in *Cyp2B6* expression in the low GGT group did not allow for statistical significance. Gene expression profile of mid GGT group revealed a similar trend to that of high GGT (Supplementary Fig. 5A). In contrast to phase I changes, most of the phase II genes remained unchanged during biliary atresia (Fig. 4B, Supplementary Fig. 5B), except for reduction in *Sult1E1* and *Ugt3A1* expression (Fig. 4B, Supplementary Fig. 5B). It is important to note that these two genes are known to be suppressed upon CAR and PXR activation (28–30), indicating that the xenosensing mechanism may be active in patients with high GGT, which represents about 25 percent of the biliary atresia patients.

**Fig. 4:**
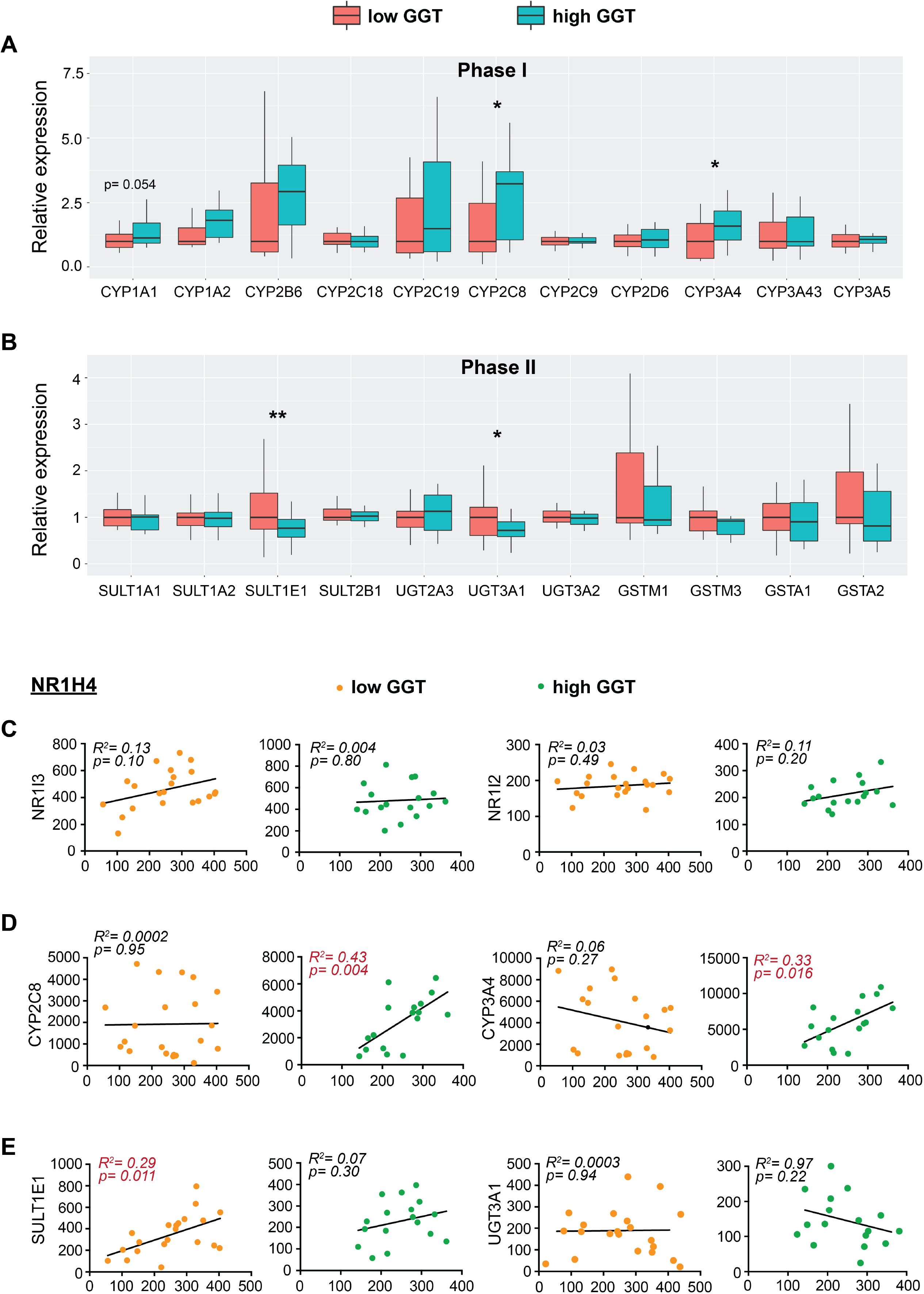
Expression and correlation with FXR mRNA levels of detoxification genes change with circulating GGT levels. Based on drug metabolic gene expression, microarray data for Biliary Atresia (GEO dataset: GSE46995) was segregated into two groups: low GGT <450 IU/l (n=21) and high GGT >850 IU/l (n=17). Box plots show relative expression for (**A**) Phase I and (**B**) Phase II genes. mRNA expression was normalized to the median value of the low GGT group (Student t test, *p <0.05). Scatter plots show the extent of correlation between FXR and (**C**) xenosensing nuclear receptors: NR1I3 (CAR) and NR1I2 (PXR), (**D**) phase I genes: CYP2C8 and CYP3A4, (**E**) phase II genes: SULT1E1 and UGT3A1. Pearson correlation coefficient test was performed. R-squared values are reported (*p <0.05).

We additionally examined if there is a correlation of *Fxr* gene expression with phase I or II genes. Interestingly, we did not see any correlation of *Fxr* expression with either *Car* or *Pxr* (Fig. 4C) but found a direct correlation of *Fxr* with Cyp3A4 as previously reported (25) and with *Cyp2C8* (Fig. 4D). This Cyp2C correlation with *Fxr* could be secondary to hepatocyte nuclear factor 4α (*Hnf4α*) activation, which is known to regulate Cyp2C8 (31). Similarly, *Sult1E1* associated with Fxr in the low GGT samples (Fig. 4E). Next, we examined correlations with *Shp* expression and observed a positive trend with CAR and PXR at the high GGT group (Supplementary Fig. 6A). *Shp* expression did not associate with *Cyp3A4 nor Cyp2C8* (Supplementary Fig. 6B), nor with *Sult1E1* nor *Ugt3A1* (Supplementary Fig. 6C), suggesting that loss of expression of SHP is not responsible for the increase in detoxification genes observed in cholestasis.

### CAR is responsible for increased phase I gene expression in cholestasis

We validated our RNA-seq data such that many of the phase I and II genes were induced in the cholestatic DKO mice (Fig. 5A). *Cyp2b10* and *Cyp3a11* are murine homologs of human *Cyp2B6* and *Cyp3A4*. It is well known that Cytochrome P450s (*Cyps*) are responsible for oxidative metabolism (32), with CYP2Bs and CYP3As carrying out phase I metabolism of many prescription drugs (Supplementary Table 2). We also analyzed Cyp2B protein expression and found its levels increased in DKO livers compared to WT (Fig. 5B). Since DKO mice displayed higher expression of Cyps, sulfotransferase *Sult2a1*, and glutathione S transferases *Gstm2* and *Gstt3*, indicating higher drug metabolizing ability, we tested its functional relevance. WT and DKO mice were treated with the paralytic agent zoxazolamine, and paralysis time was measured. Consistent with the observed increases in the detoxification machinery, DKOs showed lower paralysis time compared to WT mice (Fig. 5C). Next, we tested if this is secondary to CAR activation in the DKO animals by pretreating mice with a CAR inverse agonist Androstanol to inhibit CAR activity. We found that androstanol significantly increased paralysis time in WT indicating that CAR downstream target genes are important to clear Zoxazolamine (Fig. 5C). Notably, DKO mice also lost their increased drug clearance capacity when treated with androstanol, indicating that CAR activation in DKO mice contributes towards faster zoxazolamine removal.

**Fig. 5.**
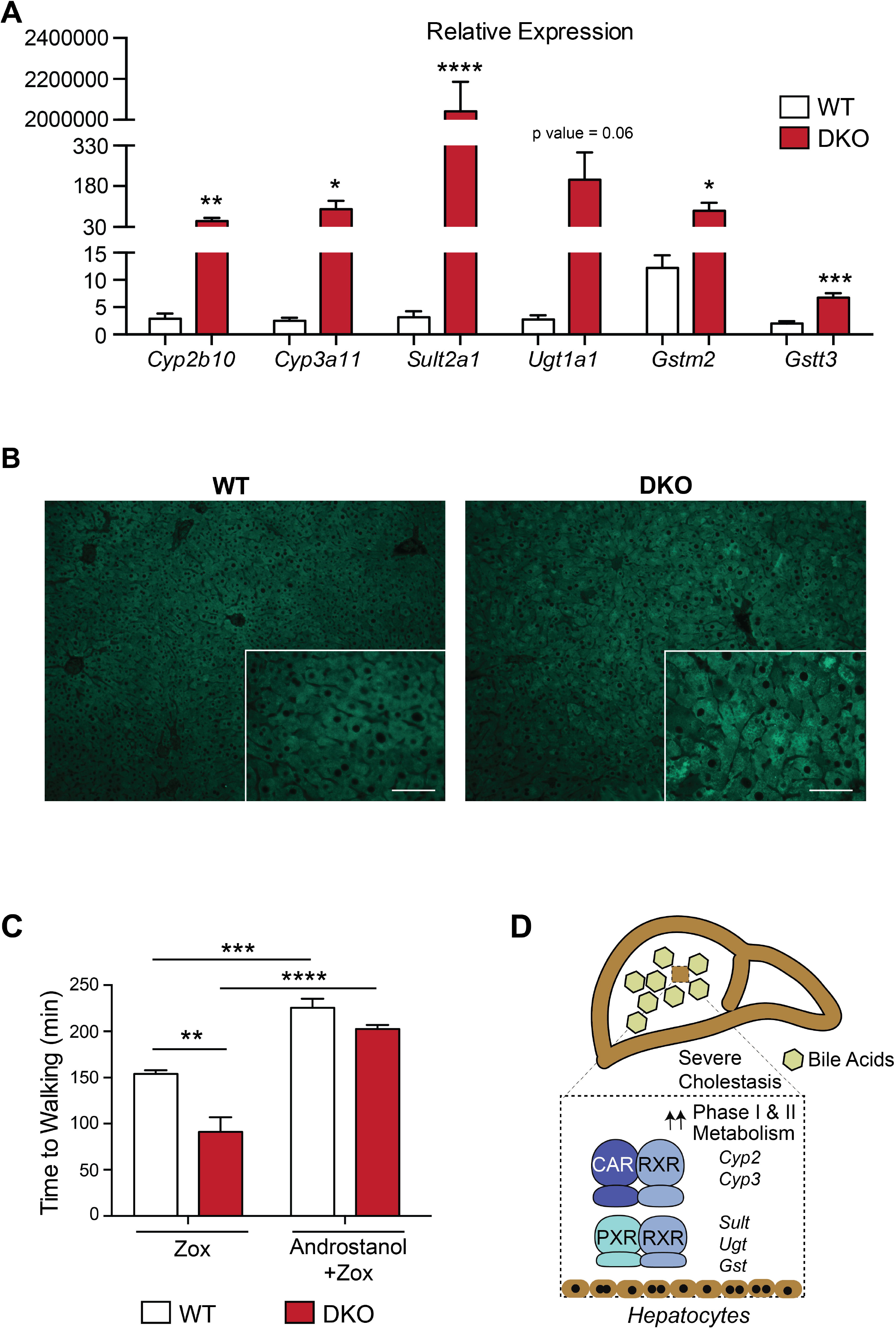
Increased drug metabolism in cholestatic mice is attributed to increased activation of CAR. (**A**) qPCR analysis shows upregulation of key genes involved in Phase I and Phase II detoxification in DKO livers (n=5 mice per group, Student t test, *p <0.05). (**B**) DKO cholestatic livers show increased immunofluorescence staining for CYP2B compared to WT livers (n=4-5 mice per group, Scale Bar: 50μ). (**C**) Bar graph indicates that DKO mice recover faster post zoxazolamine (zox)-induced paralysis than WT mice. Inhibiting CAR activity with androstanol normalizes the difference in paralysis time between WT and DKO mice (n=3-4 mice per group, mean ± SEM, One-way ANOVA, **p <0.001). (**D**) Schematic figure demonstrating CAR activation in cholestasis.

Several studies in the 1970s reported faster clearance of drugs in patients with liver disease. For instance, tolbutamide was metabolized faster in cholestatic patients (165 ± 48 min) compared to normal individuals (384 ±76 min) (33). Patients with chronic alcohol consumption had higher metabolism of antipyrine (11.7 hr vs. 15.7 hr), phenobarbital (26.3 hr vs. 35.1 hr), and warfarin (26.5 hr vs. 41.1 hr) compared to controls (12). It is important to note that many of these drugs are prototypical substrates of the CAR/PXR target genes *Cyp2C, 2B*, or *3A* (Supplementary Table 2). Taken together, our findings implicate that hepatobiliary damage can activate the CAR-PXR network, which in turn leads to unexpected increases in drug metabolizing ability (Fig. 5D).

## Discussion

Liver is a central organ responsible for drug metabolism and breakdown, therefore it can directly affect therapeutic efficacy. Many liver diseases with different etiologies exhibit varying levels of bile acid (BA) accumulation (34–36). Due to their detergent nature, BAs, when in excess, can also disrupts cell membranes, injures liver cells, and cause cell death (3). Therefore, BA concentrations are tightly controlled and in fact, BAs auto-regulate their concentrations via the nuclear receptors FXR and SHP (37, 38). In this study, we took a genome-wide approach to investigate pathways that are altered under different conditions of liver diseases using mouse models and by mining publicly available human data. We used *Fxr* and *Shp* double knockout mice which mimics both juvenile (14) and neonatal cholestasis Progressive Familial Intrahepatic Cholestasis 5 (PFIC5) (39) along with fatty and regenerating murine livers for our analysis. We were intrigued to find a specific increase in metabolic gene expression during cholestasis.

This was confounding, as one would expect diseased livers to have poor metabolic function. Therefore, we compared RNA-seq analysis of different liver injury cohorts with activation of xenosensors CAR or PXR activated groups, since it was previously shown that activating CAR and PXR could detoxify and eliminate BAs (40). We also examined if individual loss of FXR and SHP can control targets of CAR and PXR and found a poor overlap of target gene expression between these groups. Further, FXRKO as well as SHPKO do not accumulate as much BAs compared to DKO or 1% CA-diet fed mice (14), indicating that excess BAs may be required to promote this activation.

To elucidate the clinical significance of these findings, we mined the available datasets for different liver disease states due to HBV or HCV infections, alcoholic hepatitis, and biliary atresia. Humans have lesser number of P450s compared to mice, but do have a number of allelic forms (41). Despite dissimilarities, it is well established that nuclear receptors CAR and PXR regulate xenobiotic metabolism, and contribute to overall drug clearance (42, 43). In patients, we did not find increases in expression of drug metabolism genes in any of these liver disease conditions, except in biliary atresia samples with GGT levels >850U/L. While we do not know why such a significant correlation exists only in this high GGT group, it is important to note that this subset accounts for a significant 25 percent of these patients and shows induction in CAR-PXR target genes. All biliary atresia patients displayed elevated conjugated bilirubin levels to the same extent (~5 mg/dl), which unlike serum GGTs, did not separate into distinct clusters.

Even though we see high expression at the gene level, we need to validate the P450 activity in different human cholestatic conditions, which is challenging. Instead, we have validated these findings at the protein level using our cholestatic mouse model and found higher Cyp2B protein expression in DKO compared to WT mice livers.

Moreover, we tested the functionality of these changes by measuring the effect of zoxazolamine in these animals, and verified that indeed, the detoxification machinery is activated in DKO and therefore they can metabolize this drug faster.

In summary, we postulate that severely cholestatic human hepatocytes may mount detoxification response to alleviate some of the BA burden. Although this may be initiated for a beneficial purpose, it comes with adverse consequences in terms of therapeutic benefits. As *Cyp2C8, 2B6* and *3A4* are responsible for the majority of therapeutic drug clearance, these findings pose a challenge in the management of disease in this subset of patients. These individuals will have lower therapeutic benefit since they will clear prescription drugs faster than normal patients, as previously reported with certain drugs (tolubutamide, warfarin, antipyrine and phenytoin) (12, 33). Based on our study, we propose that monitoring serum GGT levels, combined with therapeutic dose adjustments will be invaluable for patients with overt cholestasis during their disease management.

## Materials and Methods

### Animals

*Fxr^−/−^ Shp^−/−^*(DKO) male mice were generated as previously described (14). Age matched wild-type (WT) males were used as controls. Individual floxed-Fxr and floxed-*Shp* mice obtained from Dr. Kristina Schoonjans’s laboratory were intercrossed to generate double floxed homozygous *Fxr-Shp* mice (fl/fl *Fxr*fl/fl *Shp*). Mice were maintained on C57BL/6J background, and housed on a standard 12-hour-light/dark cycle. All the experiments were carried out on 12-16 weeks old male mice, unless specified, as outlined in the Guide for the Care and Use of Laboratory Animals, the National Academy of Sciences (NIH publication 86-23, revised 1985) and approved by the Institutional Animal Care and Use Committee at University of Illinois, Urbana-Champaign. To test if increased detoxification is a generalized injury response of the liver, following experimental cohorts were used- (i) Bile acid-induced stress: fl/fl *Fxr* fl/fl *Shp* mice were fed with either normal chow or 1% Cholic Acid (TD.140532) diet for 5 days. (ii) High-fat diet induced fatty liver: WT mice were fed either a chow or 60% fat containing diet (TD.06414) for 8 weeks. (iii) Liver regeneration model: Livers from C3H/HeN mice were collected 48 hours post 2/3^rd^ partial hepatectomy (PHx) to determine if CAR is activated in the regenerating liver. All the mouse diets were obtained from Harlan (Envigo).

### Chemicals

Murine CAR agonist TCPOBOP (1,4-Bis [2-(3,5-dichloropyridyloxy)] benzene), referred here as TC, was obtained from Sigma. TC was first dissolved in 100% ethanol at a concentration of 4 mg/ml, followed by dilution in corn oil and stirred overnight to make a final concentration of 2 mg/ml. Thirty-six hours after a single intraperitoneal injection with corn oil or 3 mg/kg TC, WT mice were sacrificed to collect livers. WT and DKO mice were injected intraperitoneally once with either 250 mg/kg zoxazolamine (Sigma) alone, or with 100 mg/kg androstanol (Sigma), 3 days prior to zoxazolamine treatment to inhibit CAR activity.

### Histology and Immunofluorescence Staining

Liver tissues were fixed in 10% Formalin for 24 hours, processed and embedded in paraffin. Liver sections were cut to obtain 5μ thick slices, followed by deparaffinization in xylene and ethanol (100%, 95%, 80% and 50%) and rehydration in water. These slices were stained with Hematoxylin and Eosin as per manufacturer’s protocol.

For immunofluorescence staining, the rehydrated sections were antigen retrieved in Tris-EDTA buffer (pH 8.0) in a slow cooker at 90□°C for 5□min, followed by cool down in cold tap water. Next, the sections were washed in Tris-buffered saline (TBS)□+□0.05% Triton X-100 buffer twice for 5 minutes, and blocked in 10% normal goat serum and 1% BSA at room temperature for 2 hours. Post blocking, anti-CYP2B (Origene, Catalog# TA504328) were applied to the sections at 1:100 dilution and incubated overnight at 4□°C. Following this, the sections were washed in TBS+ Triton X-100 buffer, and secondary fluorescent antibodies (Goat anti-mouse, 1:500 in 1% BSA + TBS) were applied for 1□h at room temperature. The sections were then coverslipped using CC/Mount aqueous mounting media (Sigma), and imaged on Zeiss LSM 710 microscope at Institute for Genomic biology (IGB) core facility, UIUC.

### RNA-Seq analysis

RNA was isolated using RNeasy tissue mini-kit (Qiagen) from the livers of the following cohorts of mice: Cohort I: Chow fed WT and DKO (*Fxr^−/−^Shp^−/−^* double knockout) mice, Cohort II: Corn Oil- or TC-treated WT mice, Cohort III: Chow or High fat diet-fed WT mice, Cohort IV: Sham- or partial hepatectomized (PHx) mice, Cohort V: Chow- or Cholic acid (CA)-fed mice. RNA quality was tested using an Agilent bioanalyzer by the Functional Genomics Core at Roy J. Carver Biotechnology Center, UIUC. Hi-Seq libraries were prepared and Illumina sequencing was performed on a HiSeq 4000 at the High Throughput Sequencing and Genotyping Unit, UIUC. The sequencing details are outlined in Supplementary Table 3. Reads were processed for quality using Trimmomatic and aligned to the mouse genome (mm10) using STAR. Read count values and differential gene expression were obtained from HTseq and DESeq2 respectively. Fold change >1.5, calculated relative to the respective controls, was considered significant. Log2 fold change was plotted as heat maps. Upregulated and downregulated genes were clustered into biological pathways using DAVID Gene-Ontology tool (https://david.ncifcrf.gov/tools.jsp).

The data discussed in this publication have been deposited in NCBI’s Gene Expression Omnibus (44) and are accessible through GEO Series accession number GSE116627 (https://www.ncbi.nlm.nih.gov/geo/query/acc.cgi?acc=GSE116627).

### Quantitative Venn Diagrams

BioVenn (45) was used to generate Venn diagrams comparing the overlapping Phase I and Phase II metabolic genes between the following groups: Group I: DKO, CAR-activated and PXR-activated mice, Group II: 5 weeks old DKO, *Fxr^−/−^* (FXRKO), *Shp^−/−^* (SHPKO) mice (14), Group III: DKO and Cholic Acid (CA)-fed mice. Genes showing similar trend to DKO directionally were included. Difference of >25% in gene expression between the groups was considered significant.

### Analysis of Serum Bile acid composition

Pooled serum from either DKO, CA-diet fed mice or their respective controls (n=5 mice per group) was quantitated for individual bile acid species using liquid chromatographic– mass spectrometry at Metabolomics Core (Baylor College of Medicine, Houston). A Waters Acquity UPLC BEH C18 column was used. L-Zeatine was spiked into each sample as an internal standard.

### Quantitative real time PCR (qRT-PCR) Analysis

RNA was isolated from WT and DKO livers (n=5 per group) using TRIzol (Invitrogen), reverse-transcribed, and analyzed by SYBR^®^ Green™ based qRT-PCR. Mouse specific primer sets are listed in Supplementary Table 4. Relative gene expression was calculated by delta-delta Ct method, and normalized to *36b4* levels as loading control.

### Statistics

Data is presented as mean ± standard error of mean (SEM). Unpaired Student’s t test was used to compare between WT and DKO. One-way ANOVA with a Bonferroni post-hoc test was used for the zoxazolamine experiment. All statistical analysis was done using Graphpad Prism. Correlation was assessed using Pearson correlation coefficient test. Level of significance: \**p* value <0.05.

## Author contributions

B.M., W.A., and M.E.P. did the experiments and data analysis; B.M., W.A., M.E.P., A.K. and S.A. did the interpretation of data, and B.M. and S.A. wrote the manuscript; R.F. and A.W. were responsible for partial hepatectomy and regeneration studies; B.M., R.F., and W.A. performed and analyzed RNA sequencing; M.E.P. and S.A. performed the zoxazolamine experiment; M.E.P. with input from S.A. put together the drug table; B.M. and W.A. analyzed the publicly available transcriptome data. S.A. was responsible for the study design, editing the manuscript, and study supervision.

