## Supplementary Figures and legends for "Constitutive Androstane Receptor contributes towards increased drug clearance in cholestasis"

**Supplementary Figure Legends**

**Supplementary Fig. 1: DKO livers demonstrate ballooning hepatocytes, focal inflammation, ductular proliferation and necrosis.**
Hematoxylin and Eosin staining was performed on WT, DKO, HFD fed, CA fed, and PHx livers. WT section shows a normal liver histology, whereas DKO livers show hepatomegaly and various signs of injury. Inset displays ductular reaction. Black circle: necrotic patch, yellow circles: focal inflammation, white arrow: microsteatosis. As expected, HFD display fat accumulation in the liver, and CA diet and PHx livers exhibit proliferating hepatocytes. Black arrows: mitotic bodies. Scale: 100μ.

**Supplementary Fig. 2. Bile acid overload is sufficient to elicit comparable detoxification response observed in DKO livers.**
**(A)** Bar graphs show upregulated and downregulated biological pathways in bile acid-stressed livers (n=2-3 mice per group, Fold change >1.5, FDR <20). Venn Diagrams display overlapping **(B)** Phase I and **(C)** Phase II genes between DKO and CA fed mice, indicating enhanced drug clearance in DKO livers as a specific response to increased bile acid levels (n=3 per group, Fold change > 1.5, p value <0.05). **(D)** Bile acid composition analysis reveals increased 12 alpha-hydroxy species in serum from DKO and CA fed mice. Fold change values compared to the respective controls were plotted. (n=5 per group).

**Supplementary Fig. 3. Transcript profiles in DKO cholestatic livers mimic CAR activated livers.**
Venn diagrams quantitate significant similarity in **(A)** overall**, (B)** Phase I, and **(C)** Phase II gene changes between DKO and CAR activated livers. Numbers indicate total number of genes in the respective livers as well as the overlap of DKO with TC- injected mice (n=3 per group, Fold change > 1.5, p value <0.05).

**Supplementary Fig. 4. Loss of FXR mimics only Phase II metabolism in DKO livers.**
Heat maps displaying expression profiles of **(A)** Phase I and **(C)** Phase II genes in livers isolated from 5 weeks old DKO, FXRKO and SHPKO. Overlap was quantitated in Venn Diagrams **(B-D)**. Phase I genes are equally distributed between the individual FXRKO and SHPKO, compared to DKO livers. However, FXRKO livers show Phase II gene changes similar as DKO livers. No overlap between DKO and SHPKO was observed. Numbers indicate total number of genes in the respective genotypes as well as overlap of DKO with FXRKO and SHPKO (n=3 per group, Fold change > 1.5, p value <0.05).

**Supplementary Fig. 5. Differential expression of genes regulating phase I, II and steroid metabolism, and nuclear receptors in biliary atresia patients.**
Biliary atresia microarray data (GEO dataset: GSE46995) was categorized into three cohorts on the basis of their GGT levels: low (GGT< 450 IU/l, n=21), mid (450< GGT< 850 IU/l, n=33) and high (GGT >850 IU/l, n=17). Box plots show relative expression of **(A)** phase I, **(B)** phase II, **(C)** nuclear receptors, and **(D)** steroid metabolic genes. Gene expression was normalized to the median value of the low GGT group (Student t test, *p <0.05).

**Supplementary Fig. 6: SHP and detoxification gene profiles do not show significant correlations.**

SHP Correlations were compared in the biliary atresia dataset (GSE46995) between low (n= 21) and high GGT (n=17) groups. Scatter plots show the extent of correlation between SHP and **(A)** xenosensors: NR1I3 (CAR) and NR1I2 (PXR), **(B)** phase I genes CYP2C8 and CYP3A4, **(C)** phase II genes: SULT1E1 and UGT3A1 (Pearson correlation coefficient test, *p <0.05).

**Supplementary Table 1: Gene Ontology analysis of gene expression**

**Supplementary Table 2: Some common prescription drugs metabolized by CAR and PXR target.**

**Supplementary Table 3: RNA-seq experimental set up and analysis.**

**Supplementary Table 4: Primers used for qRT-PCR assays.**

**Supplementary Figure 1**

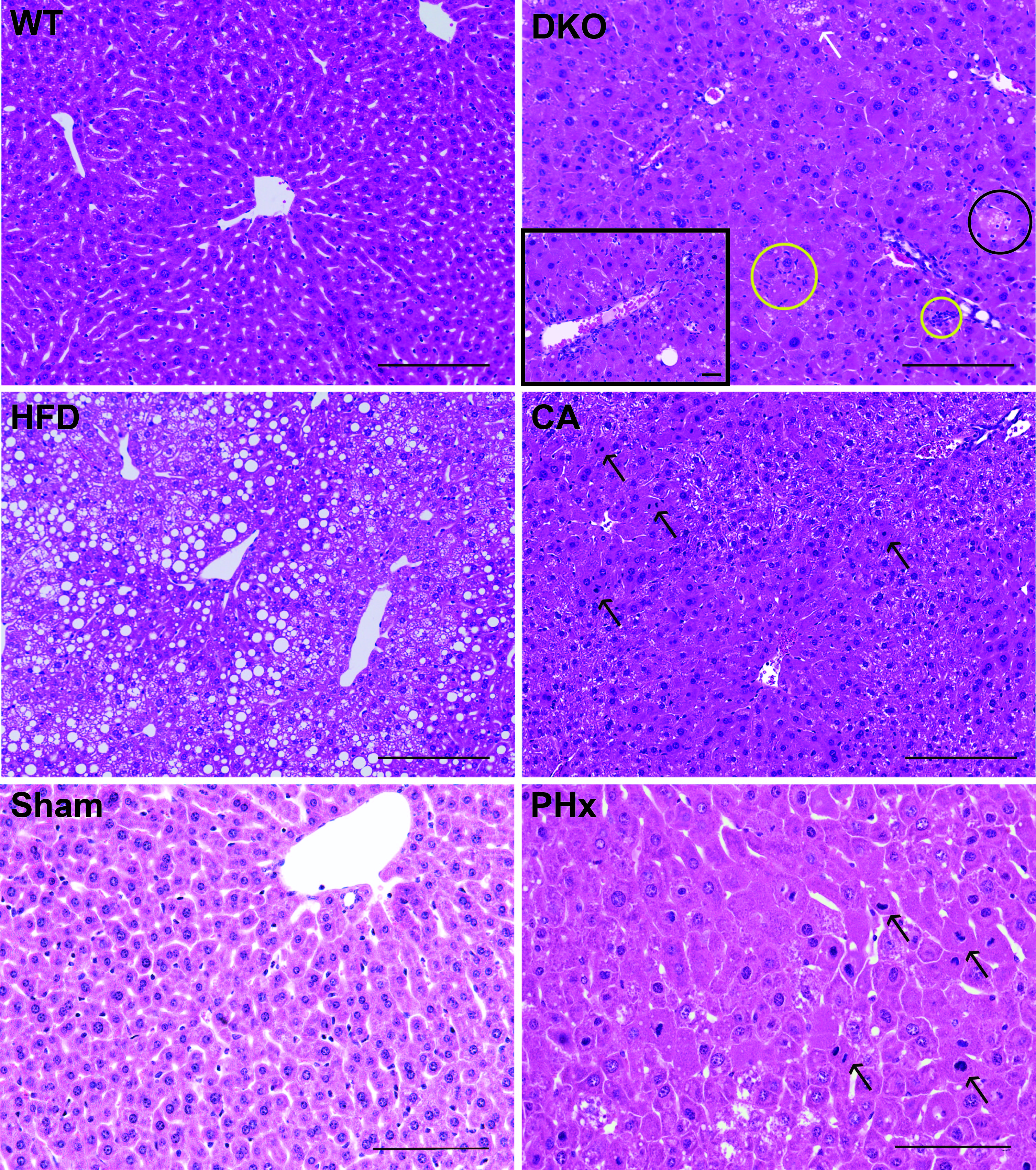

**Supplementary Figure 2**

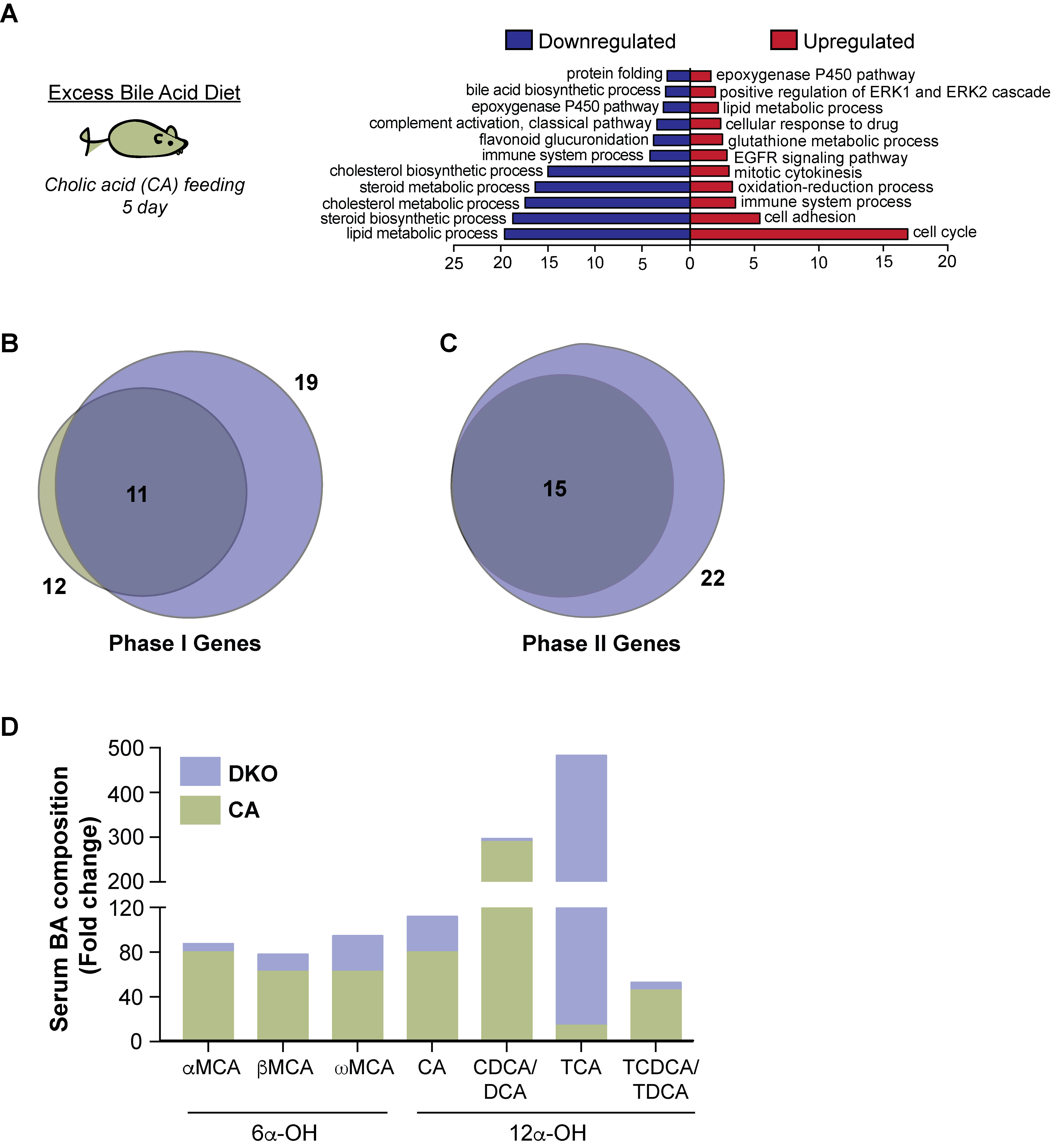

**Supplementary Figure 3**

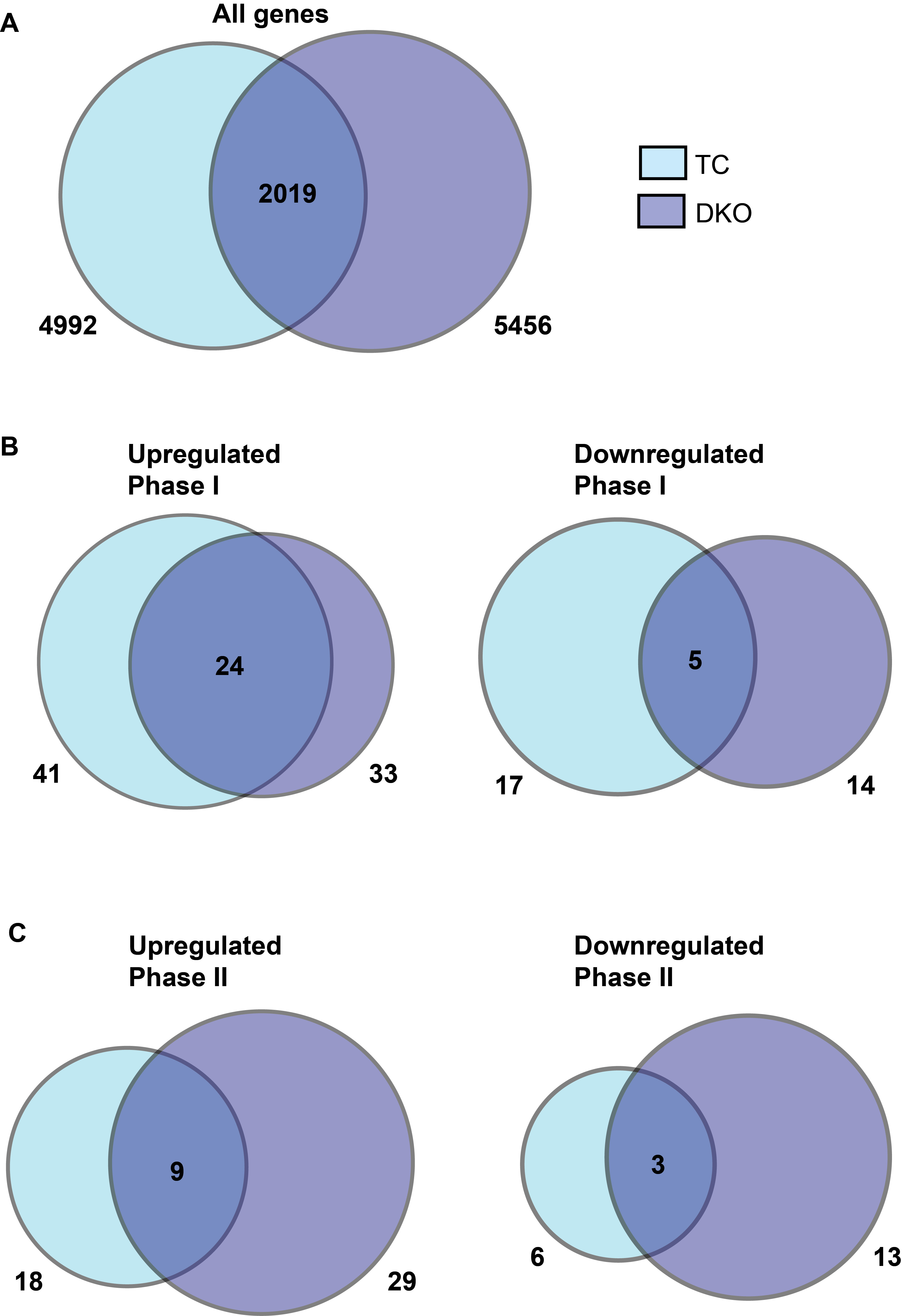

**Supplementary Figure 4**

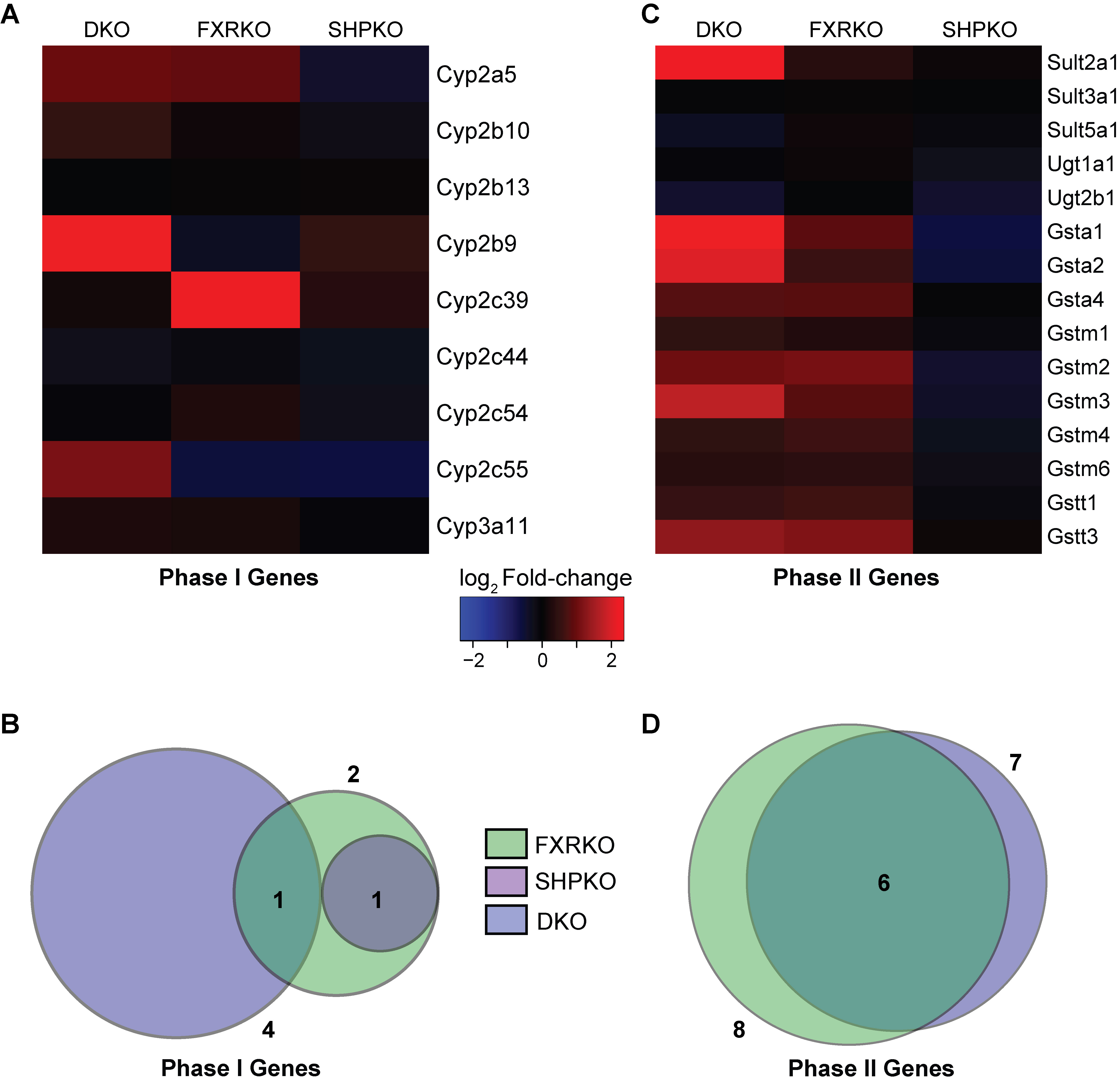

**Supplementary Figure 5**

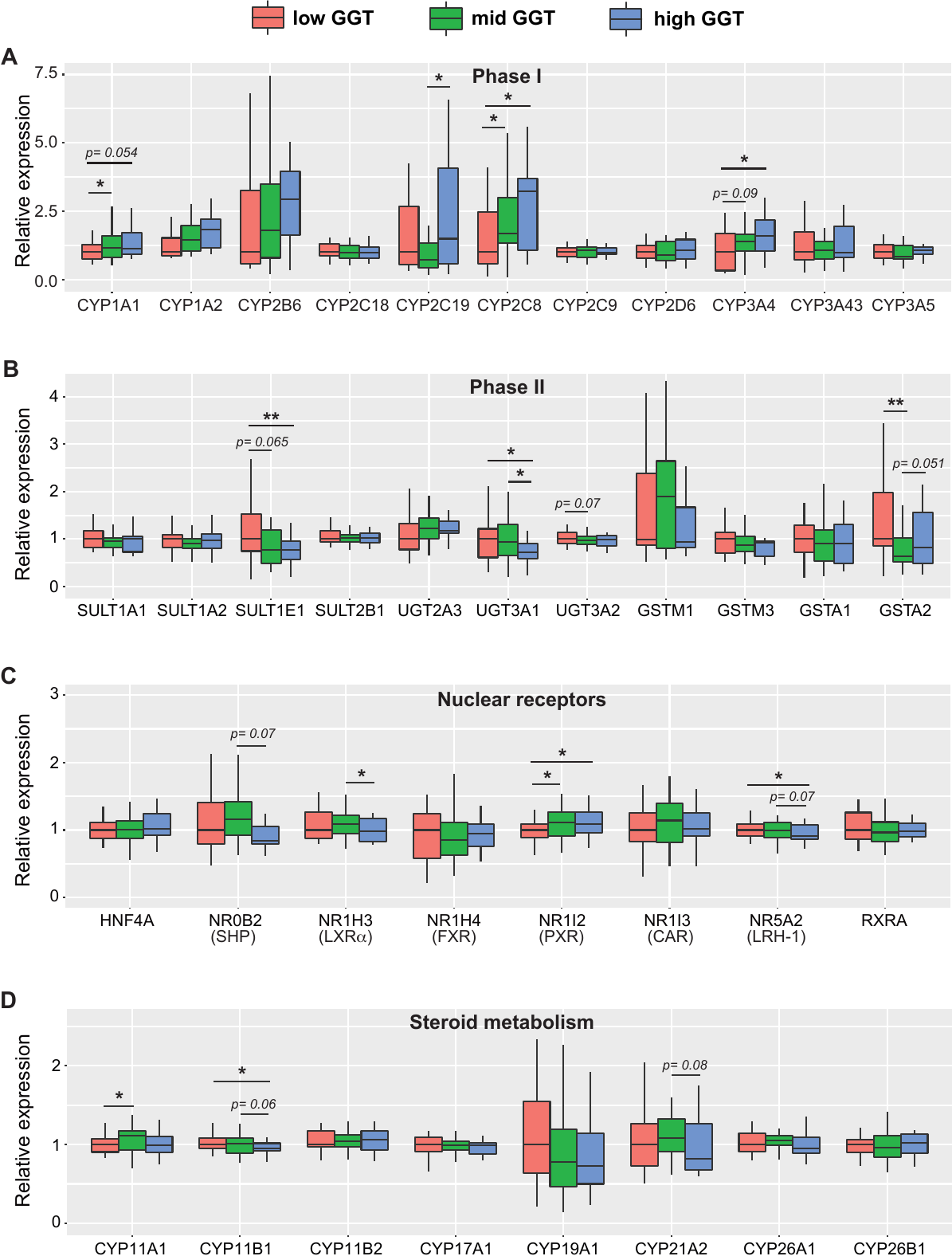

**Supplementary Figure 6**

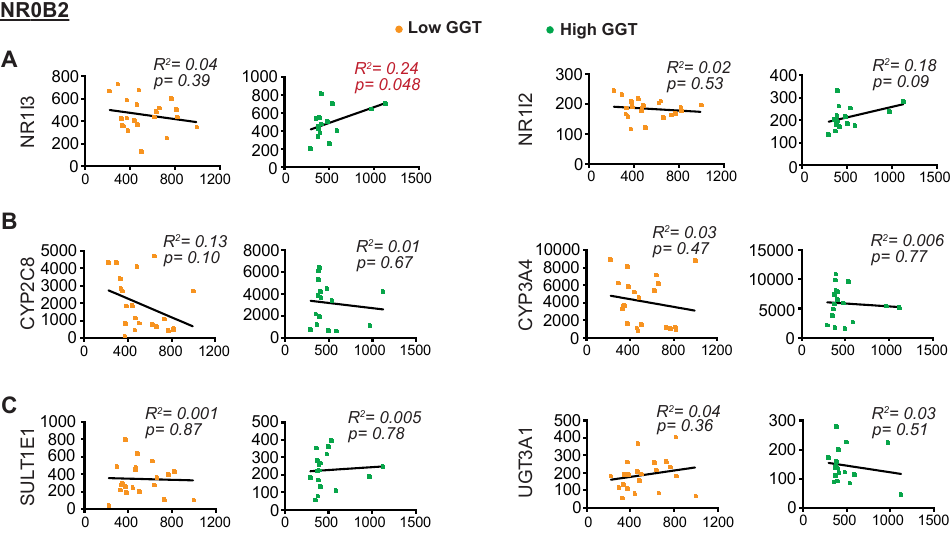
