## Supplementary table 1 for "Constitutive Androstane Receptor contributes towards increased drug clearance in cholestasis"

| Category | Term | Count | % | PValue | Genes | List Total | Pop Hits | Pop Total | Fold Enrichm | Bonferroni | Benjamini | FDR |
|---|---|---|---|---|---|---|---|---|---|---|---|---|
| GOTERM_BP | GO:0007049~cell cycle | 74 | 16.59192825 | 8.71E-36 | KIF23, E2F1, CLSP | 365 | 614 | 18082 | 5.97058587 | 1.55E-32 | 1.55E-32 | 1.48E-32 |
| GOTERM_BP | GO:0007067~mitotic nuclear division | 47 | 10.53811659 | 6.46E-29 | KIF23, LRRCC1, KI | 365 | 277 | 18082 | 8.40565748 | 1.15E-25 | 5.73E-26 | 1.09E-25 |
| GOTERM_BP | GO:0051301~cell division | 51 | 11.43497758 | 7.34E-27 | KIF23, KIFC1, PRC | 365 | 374 | 18082 | 6.75541719 | 1.30E-23 | 4.34E-24 | 1.24E-23 |
| GOTERM_BP | GO:0000070~mitotic sister chromatid segregation | 10 | 2.242152466 | 3.14E-10 | CDCA8, MAD2L1, | 365 | 23 | 18082 | 21.5390113 | 5.56E-07 | 1.39E-07 | 5.31E-07 |
| GOTERM_BP | GO:0007059~chromosome segregation | 15 | 3.3632287 | 2.73E-09 | DSN1, ESPL1, CEN | 365 | 89 | 18082 | 8.34939203 | 4.84E-06 | 9.67E-07 | 4.62E-06 |
| GOTERM_BP | GO:0007080~mitotic metaphase plate congression | 10 | 2.242152466 | 1.65E-08 | CCNB1, KIF22, KII | 365 | 34 | 18082 | 14.5705077 | 2.94E-05 | 4.89E-06 | 2.81E-05 |
| GOTERM_BP | GO:0000281~mitotic cytokinesis | 7 | 1.569506726 | 2.52E-05 | CKAP2, KIF23, PLI | 365 | 30 | 18082 | 11.5592694 | 0.04373771 | 0.00636864 | 0.0427072 |
| GOTERM_BP | GO:0007018~microtubule-based movement | 10 | 2.242152466 | 2.64E-05 | KIF23, KIF22, KIF2 | 365 | 78 | 18082 | 6.35124693 | 0.04575435 | 0.00583717 | 0.0447227 |
| GOTERM_BP | GO:0034501~protein localization to kinetochore | 5 | 1.121076233 | 3.08E-05 | CDK1, MTBP, TTK | 365 | 10 | 18082 | 24.769863 | 0.05323553 | 0.00605989 | 0.05223667 |
| GOTERM_BP | GO:0006915~apoptotic process | 27 | 6.053811659 | 9.55E-05 | E2F1, SIVA1, S10( | 365 | 570 | 18082 | 2.3466186 | 0.1559516 | 0.01681162 | 0.16180691 |
| GOTERM_BP | GO:0030199~collagen fibril organization | 7 | 1.569506726 | 1.19E-04 | LUM, COL3A1, CC | 365 | 39 | 18082 | 8.8917457 | 0.19019169 | 0.01899525 | 0.20128921 |
| GOTERM_BP | GO:0031100~organ regeneration | 8 | 1.793721973 | 1.26E-04 | PKM, LIF, CDK1, ( | 365 | 56 | 18082 | 7.07710372 | 0.19978233 | 0.01840122 | 0.21264488 |
| GOTERM_BP | GO:0006974~cellular response to DNA damage stimulus | 22 | 4.932735426 | 1.31E-04 | CLSPN, RAD51B, I | 365 | 420 | 18082 | 2.59493803 | 0.20700757 | 0.01768344 | 0.22128921 |
| GOTERM_BP | GO:0051256~mitotic spindle midzone assembly | 4 | 0.896860987 | 1.55E-04 | KIF23, KIF4, AURI | 365 | 6 | 18082 | 33.026484 | 0.24010977 | 0.01942187 | 0.26191726 |
| GOTERM_BP | GO:0007051~spindle organization | 5 | 1.121076233 | 2.43E-04 | SPAG5, AURKA, A | 365 | 16 | 18082 | 15.4811644 | 0.35003118 | 0.0283135 | 0.41065407 |
| GOTERM_BP | GO:0051988~regulation of attachment of spindle microtubules to kinetochore | 4 | 0.896860987 | 2.67E-04 | SPAG5, KNSTRN, | 365 | 7 | 18082 | 28.3084149 | 0.37709955 | 0.02915217 | 0.45110802 |
| GOTERM_BP | GO:0006935~chemotaxis | 10 | 2.242152466 | 6.40E-04 | S100A8, CCR5, S1 | 365 | 118 | 18082 | 4.19828187 | 0.67912146 | 0.06467786 | 1.07981703 |
| GOTERM_BP | GO:0006260~DNA replication | 10 | 2.242152466 | 8.63E-04 | TICRR, DTL, FAM1 | 365 | 123 | 18082 | 4.02762 | 0.78410764 | 0.08163953 | 1.45352379 |
| GOTERM_BP | GO:0006268~DNA unwinding involved in DNA replication | 4 | 0.896860987 | 8.74E-04 | MCM2, MCM4, T | 365 | 10 | 18082 | 19.8158904 | 0.78816862 | 0.07843535 | 1.47139586 |
| GOTERM_BP | GO:0000278~mitotic cell cycle | 6 | 1.34529148 | 9.35E-04 | PRDM5, KIF18B, ( | 365 | 38 | 18082 | 7.822062 | 0.80990162 | 0.07965868 | 1.57321367 |
| GOTERM_BP | GO:0006270~DNA replication initiation | 5 | 1.121076233 | 0.00124774 | MCM2, MCM3, N | 365 | 24 | 18082 | 10.3207763 | 0.89096817 | 0.1001519 | 2.09443427 |
| GOTERM_BP | GO:0050729~positive regulation of inflammatory response | 7 | 1.569506726 | 0.00165132 | S100A8, CCR5, S1 | 365 | 63 | 18082 | 5.504414 | 0.94679026 | 0.12483386 | 2.76299865 |
| GOTERM_BP | GO:0051726~regulation of cell cycle | 9 | 2.01793722 | 0.00192951 | E2F1, CDKN1A, C( | 365 | 112 | 18082 | 3.98087084 | 0.96755457 | 0.13847567 | 3.22134569 |
| GOTERM_BP | GO:0001578~microtubule bundle formation | 5 | 1.121076233 | 0.00196515 | PRC1, PLK1, PSRC | 365 | 27 | 18082 | 9.17402334 | 0.96954727 | 0.13539497 | 3.27991789 |
| GOTERM_BP | GO:0007076~mitotic chromosome condensation | 4 | 0.896860987 | 0.00199147 | NCAPG, NUSAP1, | 365 | 13 | 18082 | 15.2429926 | 0.97093994 | 0.13197574 | 3.32315211 |
| GOTERM_BP | GO:0006281~DNA repair | 16 | 3.587443946 | 0.00207164 | CLSPN, KIF22, RAI | 365 | 318 | 18082 | 2.49256483 | 0.97480151 | 0.13201064 | 3.4547201 |
| GOTERM_BP | GO:0007019~microtubule depolymerization | 4 | 0.896860987 | 0.00249702 | KIF2C, KIF18A, KII | 365 | 14 | 18082 | 14.1542074 | 0.98817751 | 0.15156447 | 4.15005654 |
| GOTERM_BP | GO:0030261~chromosome condensation | 4 | 0.896860987 | 0.00372866 | CDK1, NCAPG2, S | 365 | 16 | 18082 | 12.3849315 | 0.99868081 | 0.21086031 | 6.13683345 |
| GOTERM_BP | GO:0032467~positive regulation of cytokinesis | 5 | 1.121076233 | 0.00416791 | KIF23, KIF20B, AU | 365 | 33 | 18082 | 7.5060191 | 0.99939694 | 0.22557754 | 6.83597362 |
| GOTERM_BP | GO:0000086~G2/M transition of mitotic cell cycle | 5 | 1.121076233 | 0.00416791 | CDKN1A, NES, PLI | 365 | 33 | 18082 | 7.5060191 | 0.99939694 | 0.22557754 | 6.83597362 |
| GOTERM_BP | GO:0006954~inflammatory response | 16 | 3.587443946 | 0.00434602 | S100A8, TLR1, S1( | 365 | 344 | 18082 | 2.3041733 | 0.999561 | 0.22717296 | 7.11807405 |
| GOTERM_BP | GO:0060707~trophoblast giant cell differentiation | 4 | 0.896860987 | 0.00527382 | LIF, PLK4, PRDM1 | 365 | 18 | 18082 | 11.008828 | 0.99991609 | 0.26122923 | 8.5745738 |
| GOTERM_BP | GO:0007095~mitotic G2 DNA damage checkpoint | 4 | 0.896860987 | 0.00527382 | CDK1, FANCI, MB | 365 | 18 | 18082 | 11.008828 | 0.99991609 | 0.26122923 | 8.5745738 |
| GOTERM_BP | GO:0007094~mitotic spindle assembly checkpoint | 4 | 0.896860987 | 0.00527382 | MAD2L1, PLK1, C | 365 | 18 | 18082 | 11.008828 | 0.99991609 | 0.26122923 | 8.5745738 |
| GOTERM_BP | GO:0034080~CENP-A containing nucleosome assembly | 3 | 0.67264574 | 0.00574572 | OIP5, MIS18A, CE | 365 | 6 | 18082 | 24.769863 | 0.99996386 | 0.2735801 | 9.3071206 |
| GOTERM_BP | GO:0000910~cytokinesis | 5 | 1.121076233 | 0.00631678 | KIF23, PRC1, PLK1 | 365 | 37 | 18082 | 6.69455757 | 0.99998696 | 0.28883008 | 10.1862039 |
| GOTERM_BP | GO:0034605~cellular response to heat | 5 | 1.121076233 | 0.00762366 | CDKN1A, MKI67, | 365 | 39 | 18082 | 6.35124693 | 0.99999874 | 0.32936142 | 12.1679237 |
| GOTERM_BP | GO:0010165~response to X-ray | 4 | 0.896860987 | 0.00822135 | CDKN1A, CCND1, | 365 | 21 | 18082 | 9.43613829 | 0.99999957 | 0.34207412 | 13.0604537 |
| GOTERM_BP | GO:0010811~positive regulation of cell-substrate adhesion | 5 | 1.121076233 | 0.00989821 | SMOC2, LIMS1, FI | 365 | 42 | 18082 | 5.89758643 | 0.99999998 | 0.3876604 | 15.5192056 |
| GOTERM_BP | GO:0048146~positive regulation of fibroblast proliferation | 6 | 1.34529148 | 0.0099018 | CCNB1, E2F1, CDI | 365 | 65 | 18082 | 4.57289779 | 0.99999998 | 0.37959714 | 15.524397 |
| GOTERM_BP | GO:0008361~regulation of cell size | 4 | 0.896860987 | 0.01196794 | ATP2B2, IQGAP3, | 365 | 24 | 18082 | 8.256621 | 1 | 0.43016073 | 18.4638016 |
| GOTERM_BP | GO:0051382~kinetochore assembly | 3 | 0.67264574 | 0.01324933 | CENPW, CENPT, ( | 365 | 9 | 18082 | 16.513242 | 1 | 0.45504102 | 20.2381313 |
| GOTERM_BP | GO:0030071~regulation of mitotic metaphase/anaphase transition | 3 | 0.67264574 | 0.01324933 | PLK1, CENPE, UBI | 365 | 9 | 18082 | 16.513242 | 1 | 0.45504102 | 20.2381313 |
| GOTERM_BP | GO:0030308~negative regulation of cell growth | 8 | 1.793721973 | 0.01332711 | CDKN1A, CGREF1 | 365 | 125 | 18082 | 3.17054247 | 1 | 0.44863959 | 20.3446536 |
| GOTERM_BP | GO:0016525~negative regulation of angiogenesis | 6 | 1.34529148 | 0.01497797 | LIF, ECSCR, CX3CI | 365 | 72 | 18082 | 4.1283105 | 1 | 0.47969587 | 22.5742549 |
| GOTERM_BP | GO:0007275~multicellular organism development | 32 | 7.174887892 | 0.01685636 | GDF6, PAQR8, AR | 365 | 1029 | 18082 | 1.54059401 | 1 | 0.51249473 | 25.0396776 |
| GOTERM_BP | GO:0045666~positive regulation of neuron differentiation | 7 | 1.569506726 | 0.01785916 | CCR5, GDF6, HEYI | 365 | 103 | 18082 | 3.3667775 | 1 | 0.52472982 | 26.3254374 |
| GOTERM_BP | GO:0030336~negative regulation of cell migration | 7 | 1.569506726 | 0.02027933 | RAP2B, CCR5, IL1 | 365 | 106 | 18082 | 3.27149134 | 1 | 0.56241993 | 29.3435486 |
| GOTERM_BP | GO:0006955~immune response | 12 | 2.69058296 | 0.02262461 | LIF, TNFRSF10B, ( | 365 | 272 | 18082 | 2.18557615 | 1 | 0.59451249 | 32.1569502 |
| GOTERM_BP | GO:0051310~metaphase plate congression | 3 | 0.67264574 | 0.0233433 | KIF22, KIF2C, CEN | 365 | 12 | 18082 | 12.3849315 | 1 | 0.59804961 | 32.9977694 |
| GOTERM_BP | GO:0071479~cellular response to ionizing radiation | 4 | 0.896860987 | 0.02397664 | CDKN1A, RAD51A | 365 | 31 | 18082 | 6.39222271 | 1 | 0.60009687 | 33.7305959 |
| GOTERM_BP | GO:0012501~programmed cell death | 4 | 0.896860987 | 0.02397664 | PKM, DFNA5, EDA | 365 | 31 | 18082 | 6.39222271 | 1 | 0.60009687 | 33.7305959 |
| GOTERM_BP | GO:0051297~centrosome organization | 4 | 0.896860987 | 0.02607206 | PLK1, AURKA, HA | 365 | 32 | 18082 | 6.19246575 | 1 | 0.62352904 | 36.1017732 |
| GOTERM_BP | GO:0061053~somite development | 3 | 0.67264574 | 0.03134685 | RAD51B, WNT11, | 365 | 14 | 18082 | 10.6156556 | 1 | 0.68453334 | 41.7221763 |
| GOTERM_BP | GO:0045087~innate immune response | 15 | 3.3632287 | 0.03147988 | S100A8, TLR1, S1( | 365 | 400 | 18082 | 1.85773973 | 1 | 0.67873974 | 41.8577101 |
| GOTERM_BP | GO:0030073~insulin secretion | 4 | 0.896860987 | 0.03292693 | MYO5A, IL1RN, A | 365 | 35 | 18082 | 5.66168297 | 1 | 0.68816395 | 43.3130327 |
| GOTERM_BP | GO:0050727~regulation of inflammatory response | 5 | 1.121076233 | 0.03435317 | S100A8, S100A9, | 365 | 61 | 18082 | 4.06063328 | 1 | 0.69676515 | 44.7138387 |

| | | | | | | | | | | | | |
|---|---|---|---|---|---|---|---|---|---|---|---|---|
| GOTERM_BP | GO:0000082~G1/S transition of mitotic cell cycle | 5 | 1.121076233 | 0.0361607 | CDKN1A, CCND1, | 365 | 62 | 18082 | 3.9951392 | 1 | 0.70872315 | 46.4423498 |
| GOTERM_BP | GO:0007155~cell adhesion | 17 | 3.811659193 | 0.0363779 | FLRT2, HAPLN1, ( | 365 | 485 | 18082 | 1.73644401 | 1 | 0.7041926 | 46.6465904 |
| GOTERM_BP | GO:0070374~positive regulation of ERK1 and ERK2 cascade | 9 | 2.01793722 | 0.03675327 | GLIPR2, BMPER, ( | 365 | 188 | 18082 | 2.37158263 | 1 | 0.70134745 | 46.9978488 |
| GOTERM_BP | GO:0006284~base-excision repair | 4 | 0.896860987 | 0.03796617 | MUTYH, NEIL3, M | 365 | 37 | 18082 | 5.35564606 | 1 | 0.70677972 | 48.1180229 |
| GOTERM_BP | GO:0070488~neutrophil aggregation | 2 | 0.448430493 | 0.03985689 | S100A8, S100A9 | 365 | 2 | 18082 | 49.539726 | 1 | 0.71820327 | 49.8198645 |
| GOTERM_BP | GO:0000915~actomyosin contractile ring assembly | 2 | 0.448430493 | 0.03985689 | KIF23, RACGAP1 | 365 | 2 | 18082 | 49.539726 | 1 | 0.71820327 | 49.8198645 |
| GOTERM_BP | GO:0045860~positive regulation of protein kinase activity | 5 | 1.121076233 | 0.03993771 | CDKN1A, LRP8, CI | 365 | 64 | 18082 | 3.8702911 | 1 | 0.71272295 | 49.8914318 |
| GOTERM_BP | GO:2000353~positive regulation of endothelial cell apoptotic process | 3 | 0.67264574 | 0.04026318 | AKR1C18, CD248, | 365 | 16 | 18082 | 9.28869863 | 1 | 0.70956285 | 50.1786475 |
| GOTERM_BP | GO:0051276~chromosome organization | 4 | 0.896860987 | 0.0406247 | CDCA8, CENPW, ( | 365 | 38 | 18082 | 5.214708 | 1 | 0.70680216 | 50.4958611 |
| GOTERM_BP | GO:0016310~phosphorylation | 20 | 4.484304933 | 0.04161613 | CDK1, PFKFB4, LII | 365 | 612 | 18082 | 1.6189453 | 1 | 0.70971165 | 51.3560602 |
| GOTERM_BP | GO:0018108~peptidyl-tyrosine phosphorylation | 5 | 1.121076233 | 0.04190728 | ABI2, TTK, JAK3, I | 365 | 65 | 18082 | 3.81074816 | 1 | 0.70642716 | 51.6059893 |
| GOTERM_BP | GO:0006468~protein phosphorylation | 19 | 4.260089686 | 0.04365145 | CDK1, LIMK1, STK | 365 | 576 | 18082 | 1.63412291 | 1 | 0.71563956 | 53.0780976 |
| GOTERM_BP | GO:0007250~activation of NF-kappaB-inducing kinase activity | 3 | 0.67264574 | 0.04503684 | CHIL1, TLR1, TLR4 | 365 | 17 | 18082 | 8.74230459 | 1 | 0.72142551 | 54.2172461 |
| GOTERM_BP | GO:0002224~toll-like receptor signaling pathway | 3 | 0.67264574 | 0.04503684 | TLR1, TLR4, CD18 | 365 | 17 | 18082 | 8.74230459 | 1 | 0.72142551 | 54.2172461 |
| GOTERM_BP | GO:0014002~astrocyte development | 3 | 0.67264574 | 0.04503684 | S100A8, S100A9, | 365 | 17 | 18082 | 8.74230459 | 1 | 0.72142551 | 54.2172461 |
| GOTERM_BP | GO:0006919~activation of cysteine-type endopeptidase activity involved in apoptotic proce: | 5 | 1.121076233 | 0.0460085 | TNFRSF10B, S100 | 365 | 67 | 18082 | 3.69699448 | 1 | 0.723683 | 55.0006214 |
| GOTERM_BP | GO:0032496~response to lipopolysaccharide | 9 | 2.01793722 | 0.04639596 | THBD, TNFRSF10I | 365 | 197 | 18082 | 2.26323621 | 1 | 0.72130677 | 55.3094687 |
| GOTERM_BP | GO:0002376~immune system process | 14 | 3.139013453 | 0.04716823 | S100A8, CTPS, TL | 365 | 383 | 18082 | 1.8108516 | 1 | 0.72197319 | 55.9191072 |
| GOTERM_BP | GO:0097191~extrinsic apoptotic signaling pathway | 4 | 0.896860987 | 0.04914595 | SIVA1, TNFRSF10I | 365 | 41 | 18082 | 4.833144 | 1 | 0.73164644 | 57.4448622 |
| GOTERM_BP | GO:0001774~microglial cell activation | 3 | 0.67264574 | 0.05000722 | TLR1, CX3CR1, TL | 365 | 18 | 18082 | 8.256621 | 1 | 0.73278372 | 58.0936488 |
| GOTERM_BP | GO:0050731~positive regulation of peptidyl-tyrosine phosphorylation | 6 | 1.34529148 | 0.05332302 | LIF, TLR4, LRP8, N | 365 | 101 | 18082 | 2.94295402 | 1 | 0.75080116 | 60.5055534 |
| GOTERM_BP | GO:0090002~establishment of protein localization to plasma membrane | 4 | 0.896860987 | 0.05527158 | PKP3, TSPAN5, S1 | 365 | 43 | 18082 | 4.60834661 | 1 | 0.75863491 | 61.8613448 |
| GOTERM_BP | GO:0045184~establishment of protein localization | 4 | 0.896860987 | 0.0584645 | LIMS1, IKZF1, PLK | 365 | 44 | 18082 | 4.50361146 | 1 | 0.77353452 | 63.9886885 |
| GOTERM_BP | GO:0051496~positive regulation of stress fiber assembly | 4 | 0.896860987 | 0.0584645 | LIMK1, S100A10, | 365 | 44 | 18082 | 4.50361146 | 1 | 0.77353452 | 63.9886885 |
| GOTERM_BP | GO:0031577~spindle checkpoint | 2 | 0.448430493 | 0.0591872 | SPDL1, AURKB | 365 | 3 | 18082 | 33.026484 | 1 | 0.77315516 | 64.4544535 |
| GOTERM_BP | GO:0051383~kinetochore organization | 2 | 0.448430493 | 0.0591872 | SMC2, CENPH | 365 | 3 | 18082 | 33.026484 | 1 | 0.77315516 | 64.4544535 |
| GOTERM_BP | GO:0002282~microglial cell activation involved in immune response | 2 | 0.448430493 | 0.0591872 | CX3CR1, IL33 | 365 | 3 | 18082 | 33.026484 | 1 | 0.77315516 | 64.4544535 |
| GOTERM_BP | GO:0007088~regulation of mitotic nuclear division | 3 | 0.67264574 | 0.06049921 | MKI67, KIF20B, C | 365 | 20 | 18082 | 7.4309589 | 1 | 0.77618056 | 65.2855596 |
| GOTERM_BP | GO:0033198~response to ATP | 3 | 0.67264574 | 0.06049921 | P2RX5, SLC8A1, P | 365 | 20 | 18082 | 7.4309589 | 1 | 0.77618056 | 65.2855596 |
| GOTERM_BP | GO:0010628~positive regulation of gene expression | 14 | 3.139013453 | 0.06150024 | E2F1, CDK1, LIMS | 365 | 399 | 18082 | 1.738236 | 1 | 0.77735725 | 65.907335 |
| GOTERM_BP | GO:0001501~skeletal system development | 6 | 1.34529148 | 0.06508693 | HAPLN1, COL3A1 | 365 | 107 | 18082 | 2.77792856 | 1 | 0.79233925 | 68.0501877 |
| GOTERM_BP | GO:0031572~G2 DNA damage checkpoint | 3 | 0.67264574 | 0.06600215 | CLSPN, PLK1, DTL | 365 | 21 | 18082 | 7.07710372 | 1 | 0.79278809 | 68.576319 |
| GOTERM_BP | GO:0016337~single organismal cell-cell adhesion | 6 | 1.34529148 | 0.06717799 | LIMS1, SRPX2, KII | 365 | 108 | 18082 | 2.752207 | 1 | 0.79453932 | 69.2403173 |
| GOTERM_BP | GO:0050718~positive regulation of interleukin-1 beta secretion | 3 | 0.67264574 | 0.07166442 | CCR5, PANX1, TRI | 365 | 22 | 18082 | 6.75541719 | 1 | 0.81190189 | 71.6544599 |
| GOTERM_BP | GO:1900182~positive regulation of protein localization to nucleus | 3 | 0.67264574 | 0.07747742 | LIF, CDK1, NGFR | 365 | 23 | 18082 | 6.46170339 | 1 | 0.8329189 | 74.5178788 |
| GOTERM_BP | GO:0010243~response to organonitrogen compound | 3 | 0.67264574 | 0.07747742 | CDK1, CDKN1A, C | 365 | 23 | 18082 | 6.46170339 | 1 | 0.8329189 | 74.5178788 |
| GOTERM_BP | GO:0032497~detection of lipopolysaccharide | 2 | 0.448430493 | 0.07812939 | TLR4, TREM2 | 365 | 4 | 18082 | 24.769863 | 1 | 0.83181296 | 74.8214803 |
| GOTERM_BP | GO:1901660~calcium ion export | 2 | 0.448430493 | 0.07812939 | ATP2B2, SLC8A1 | 365 | 4 | 18082 | 24.769863 | 1 | 0.83181296 | 74.8214803 |
| GOTERM_BP | GO:0002793~positive regulation of peptide secretion | 2 | 0.448430493 | 0.07812939 | S100A8, S100A9 | 365 | 4 | 18082 | 24.769863 | 1 | 0.83181296 | 74.8214803 |
| GOTERM_BP | GO:0006651~diacylglycerol biosynthetic process | 2 | 0.448430493 | 0.07812939 | MOGAT2, MOGA' | 365 | 4 | 18082 | 24.769863 | 1 | 0.83181296 | 74.8214803 |
| GOTERM_BP | GO:0051987~positive regulation of attachment of spindle microtubules to kinetochore | 2 | 0.448430493 | 0.07812939 | CCNB1, CENPE | 365 | 4 | 18082 | 24.769863 | 1 | 0.83181296 | 74.8214803 |
| GOTERM_BP | GO:0001649~osteoblast differentiation | 6 | 1.34529148 | 0.07818671 | FIGNL1, MRC2, W | 365 | 113 | 18082 | 2.63042793 | 1 | 0.82834775 | 74.8480103 |
| GOTERM_BP | GO:0007050~cell cycle arrest | 5 | 1.121076233 | 0.08066574 | CDKN1A, CGREF1 | 365 | 81 | 18082 | 3.05800778 | 1 | 0.83447642 | 75.9704912 |
| GOTERM_BP | GO:0043123~positive regulation of I-kappaB kinase/NF-kappaB signaling | 7 | 1.569506726 | 0.08097335 | LGALS1, TSPAN6, | 365 | 149 | 18082 | 2.32736968 | 1 | 0.8320856 | 76.1064415 |
| GOTERM_BP | GO:0034614~cellular response to reactive oxygen species | 3 | 0.67264574 | 0.08343282 | SLC8A1, AKR1C1ε | 365 | 24 | 18082 | 6.19246575 | 1 | 0.83785574 | 77.1676719 |
| GOTERM_BP | GO:0042127~regulation of cell proliferation | 9 | 2.01793722 | 0.08902268 | TNFRSF10B, FA2H | 365 | 227 | 18082 | 1.96413011 | 1 | 0.85403292 | 79.4169832 |
| GOTERM_BP | GO:0035634~response to stilbenoid | 3 | 0.67264574 | 0.08952259 | GSTA1, GSTA2, LY | 365 | 25 | 18082 | 5.94476712 | 1 | 0.85243017 | 79.6076372 |
| GOTERM_BP | GO:0048844~artery morphogenesis | 3 | 0.67264574 | 0.08952259 | HOXA1, WNT11, I | 365 | 25 | 18082 | 5.94476712 | 1 | 0.85243017 | 79.6076372 |
| GOTERM_BP | GO:0010800~positive regulation of peptidyl-threonine phosphorylation | 3 | 0.67264574 | 0.08952259 | CHIL1, PLK1, TRIN | 365 | 25 | 18082 | 5.94476712 | 1 | 0.85243017 | 79.6076372 |
| GOTERM_BP | GO:0009968~negative regulation of signal transduction | 4 | 0.896860987 | 0.09086532 | ARRB1, RGS19, D | 365 | 53 | 18082 | 3.73884725 | 1 | 0.85360965 | 80.1115433 |
| GOTERM_BP | GO:0016477~cell migration | 8 | 1.793721973 | 0.09199999 | NCK2, ARC, CD24 | 365 | 191 | 18082 | 2.07496235 | 1 | 0.85409445 | 80.5282041 |
| GOTERM_BP | GO:0008630~intrinsic apoptotic signaling pathway in response to DNA damage | 4 | 0.896860987 | 0.09484574 | E2F1, CRIP1, BCL2 | 365 | 54 | 18082 | 3.66960934 | 1 | 0.85988859 | 81.5373512 |
| GOTERM_BP | GO:0016321~female meiosis chromosome segregation | 2 | 0.448430493 | 0.09669123 | PLK1, TTK | 365 | 5 | 18082 | 19.8158904 | 1 | 0.86241732 | 82.1652584 |
| GOTERM_BP | GO:0010994~free ubiquitin chain polymerization | 2 | 0.448430493 | 0.09669123 | TRIM6, UBE2C | 365 | 5 | 18082 | 19.8158904 | 1 | 0.86241732 | 82.1652584 |
| GOTERM_BP | GO:0002322~B cell proliferation involved in immune response | 2 | 0.448430493 | 0.09669123 | TLR4, CD180 | 365 | 5 | 18082 | 19.8158904 | 1 | 0.86241732 | 82.1652584 |
| GOTERM_BP | GO:0045143~homologous chromosome segregation | 2 | 0.448430493 | 0.09669123 | PLK1, ESPL1 | 365 | 5 | 18082 | 19.8158904 | 1 | 0.86241732 | 82.1652584 |
| GOTERM_BP | GO:0018119~peptidyl-cysteine S-nitrosylation | 2 | 0.448430493 | 0.09669123 | S100A8, S100A9 | 365 | 5 | 18082 | 19.8158904 | 1 | 0.86241732 | 82.1652584 |
| GOTERM_BP | GO:0070098~chemokine-mediated signaling pathway | 4 | 0.896860987 | 0.09889647 | ACKR3, CXCL11, ( | 365 | 55 | 18082 | 3.60288917 | 1 | 0.86589454 | 82.8892132 |
