## Supplementary table 2 for "Constitutive Androstane Receptor contributes towards increased drug clearance in cholestasis"

| Drug Class: | Drug: | Enzyme: | Induced by: | Reference: |
|---|---|---|---|---|
| **Allergy Drugs** | loratadine, fluticasone | Cyp3a4 | PXR | Yumibe N et al., *Int Arch Allergy Immunol* , 1995. |
| **Steroids** | estrone, testosterone | Cyp2b6 | CAR, PXR | Ohe T et al., *Drug Metab Dispo,* 2000, Imaoka S et al., Biochem Pharmacol, 1996. |
| **Statins** | lovastatin, atorvastatin | Cyp3a4 | PXR | Jacobsen W et al., *Drug Metab Dispo* , 1999, Feidt DM et al., Drug Metab Dispo, 2010. |
| **Proton Pump Inhibitors** | dexlansoprazole, pantaprazole | Cyp2c19, Cyp3a4 | CAR, PXR | Vakily M et al, *Clin Drug Investig* , 2009. |
| **Anti-diabetics** | glimepiride | Cyp2c9 | CAR | Niemi M et al., *Clin Pharmacol Ther* , 2001. |
| | glyburide | Cyp2c9, Cyp3a4 | CAR, PXR | Niemi M et al., *Clin Pharmacol Ther* , 2001. |
| **Anticoagulants** | warfarin | Cyp2a6, Cyp2c9 | CAR | Miles JS, et al., *Biochem J* , 1990. |
| | phenprocoumon | Cyp2a6 | CAR, PXR | Miles JS, et al., *Biochem J* , 1990. |
| **Anesthetics/Analgesics** | ketamine | Cyp2b6 | CAR, PXR | Yanagihara Y et al., *Drug Metab Dispo* , 2001. |
| | propofol | Cyp2b6 | CAR, PXR | Oda Y et al., *Br J Clin Pharmacol* , 2001. |
| | fentanyl | Cyp3a4 | PXR | Tateishi T et al., *Anesth Analg* , 1996. |
| **Antihypertensives** | irbesartan | Cyp2c9 | CAR | Bourrié M et al., *Drug Metab Dispo* , 1999. |
| | losartan | Cyp2c9, Cyp3a4 | CAR, PXR | Yasar U et al., *Drug Metab Dispo* , 2001. |
| **Infectious Disease Drugs** | artemisinin | Cyp2b6 | CAR, PXR | Svensson US and Ashton M, *Br J Clin Pharmacol* , 1999. |
| | quinine | Cyp2c19, Cyp3a4 | CAR, PXR | Zhao XJ et al., *J Pharmacol Exp Ther* , 1996. |
| | cyclosporine | Cyp3a4 | PXR | Relling MV et al., Mol Pharmacol, 1994. |
| **Oncological Drugs** | tacrolimus | Cyp3a4 | PXR | Shiraga T et al., *Biochem Pharmacol* , 1994. |
| | lapatinib | Cyp2c19, Cyp3a4 | PXR | Nelson MH and Dolder CR, *Ther Clin Risk Manag* , 2007. |
| | sorafenib | Cyp3a4 | PXR | Ghassabian S et al., *Biochem Pharmacol* , 2012. |
| | tamoxifen | Cyp2b6 | CAR, PXR | Coller JK, et al., *Br J Clin Pharmacol* , 2002. |
