## Supplementary table 3 for "Constitutive Androstane Receptor contributes towards increased drug clearance in cholestasis"

| Experiment | TCPOBOP Treatment | | | | *Fxr*$^{-/-}$ and *Shp*$^{-/-}$ Double Knock-out (DKO) | | | | | | Partial Hepatectomy (PHx) | | | | | | High Fat Diet | | | | Cholic Acid Diet | | | | | |
|---|---|---|---|---|---|---|---|---|---|---|---|---|---|---|---|---|---|---|---|---|---|---|---|---|---|---|
| Tissue Type | Whole Liver | | | | Whole Liver | | | | | | Whole Liver | | | | | | Hepatocytes | | | | Whole Liver | | | | | |
| Age of Mice | 8 weeks | | | | 12 weeks | | | | | | 8-10 weeks | | | | | | 16 weeks | | | | 10-12 weeks | | | | | |
| Background of Mice | C57BL/6J | | | | C57BL/6J | | | | | | C3H/HeN | | | | | | C57BL/6J | | | | ff*Fxr* ff*Shp* C57BL/6J | | | | | |
| Fed State | Ad libitum | | | | Ad libitum | | | | | | Ad libitum | | | | | | 10 Hour Fast | | | | 8-10 hour fast | | | | | |
| Read Type | Paired-end | | | | Single-end | | | | | | Paired-end | | | | | | Paired-end | | | | Paired-end | | | | | |
| Read Length | 100 bp | | | | 100 bp | | | | | | 100 bp | | | | | | 100 bp | | | | 100 bp | | | | | |
| Group | Corn Oil Injection | | TCPOBOP Injection | | Wild-type C57BL/6J | | | DKO | | | 0 Hour Sham | | | 48 Hour Post PHx | | | Normal Diet | | 60% High Fat Diet | | Normal Diet | | | Cholic Acid Diet | | |
| Replicates | 1 | 2 | 1 | 2 | 1 | 2 | 3 | 1 | 2 | 3 | 1 | 2 | 3 | 1 | 2 | 3 | 1 | 2 | 1 | 2 | 1 | 2 | 3 | 1 | 2 | 3 |
| Number of Reads | 2.2E+07 | 2.2E+07 | 2.4E+07 | 2.1E+07 | 2.9E+07 | 2.4E+07 | 2.6E+07 | 2.5E+07 | 2.5E+07 | 2.9E+07 | 4.2E+07 | 4.6E+07 | 4.1E+07 | 4.1E+07 | 4E+07 | 4.4E+07 | 8E+07 | 9.2E+07 | 8.8E+07 | 8.7E+07 | 1.9E+07 | 2.2E+07 | 2.3E+07 | 2.4E+07 | 2.3E+07 | 2.4E+07 |
| Uniquely Mapped Reads | 1.8E+07 | 1.9E+07 | 1.9E+07 | 1.8E+07 | 2.2E+07 | 1.8E+07 | 2E+07 | 2.2E+07 | 2.2E+07 | 2.5E+07 | 3.5E+07 | 3.9E+07 | 3.5E+07 | 3.6E+07 | 3.4E+07 | 3.8E+07 | 6.4E+07 | 7.3E+07 | 7.8E+07 | 7.7E+07 | 1.5E+07 | 1.7E+07 | 1.7E+07 | 2E+07 | 2E+07 | 2.1E+07 |
| Mapping Rate | 82.14% | 83.58% | 79.41% | 82.45% | 75.39% | 74.76% | 75.72% | 86.48% | 85.96% | 87.07% | 84.06% | 84.75% | 84.27% | 86.27% | 85.20% | 86.77% | 80.00% | 79.44% | 88.35% | 88.61% | 78.63% | 79.06% | 74.88% | 86.27% | 87.91% | 86.97% |
| Average Mapping Rate | 82.86% | | 80.93% | | 75.29% | | | 86.50% | | | 84.36% | | | 86.08% | | | 79.72% | | 88.48% | | 77.52% | | | 87.05% | | |
