## Supplementary table 4 for "Constitutive Androstane Receptor contributes towards increased drug clearance in cholestasis"

| **Gene** | **Forward Primer (5'-3')** | **Reverse Primer (5'-3')** |
| --- | --- | --- |
| *36b4* | AGATGCAGCAGATCCGCAT | GTTCTTGCCCATCAGCACC |
| *Cyp2b10* | TTAGTGGAGGAACTGCGGAAA | CGCAAGAACTGACGGTCTG |
| *Cyp3a11* | CAGCTTGGTGCTCCTCTACC | CTCTGGGTCTGTGACAGCAA |
| *Sult2a1* | GGAACGAACTGGCTGATTGA | AGAGATGGATGGGAAGATGG |
| *Ugt1a1* | ATGGCTTTCTTCTCCGGAAT | TCAGAAAAAGCCCCTATCCC |
| *Gstm2* | TGGAATACACAGACACAAGC | CATCAATCAAGTAGGGCAGA |
| *Gstt3* | TATTTCTGTGGCTGACTTGG | CACTTCGGCTTCTACTCTCT |
